# RNAP promoter search and transcription kinetics in live *E. coli* cells

**DOI:** 10.1101/2022.12.31.522404

**Authors:** Kelsey Bettridge, Frances E. Harris, Nicolás Yehya, Jie Xiao

**Author notes:** These authors contributed equally to this work (Kelsey Bettridge, Frances E. Harris). corresponding author Jie Xiao.

## Abstract

Bacterial transcription has been studied extensively *in vitro*, which has provided indepth insight regarding transcription mechanisms. However, the live cell environment may impose different rules on transcription than the homogenous and simplified *in vitro* environment. How an RNA polymerase (RNAP) molecule searches rapidly through the vast nonspecific chromosomal DNA in the three-dimensional nucleoid space and binds a specific promoter sequence remains elusive. The kinetics of transcription *in vivo* could also be impacted by specific cellular environments including nucleoid organization and nutrient availability. In this work, we investigated the promoter search dynamics and transcription kinetics of RNAP in live *E. coli* cells. Using single-molecule tracking (SMT) and fluorescence recovery after photobleaching (FRAP) and combining with different genetic, drug inhibition, and growth conditions, we observed that RNAP’s promoter search is facilitated by nonspecific DNA interactions and largely independent of nucleoid organization, growth condition, transcription activity, or promoter classes. RNAP’s transcription kinetics, however, is sensitive to these conditions and mainly modulated at the levels of actively engaged RNAP and the promoter escape rate. Our work establishes a foundation for further mechanistic studies of bacterial transcription in live cells.

## Introduction

Transcription regulation in response to cellular and environmental stimuli must be rapid for survival, and hence, it is fundamentally a kinetic problem for any organism. An early, potentially rate-limiting step of transcription is promoter search, where RNA polymerase (RNAP) randomly samples the chromosome until it finds a promoter(Feklistov, 2013). During promoter search, RNAP must balance specificity and efficiency to allow rapid discrimination of specific promoter DNA sequences against the excess nonspecific chromosomal DNA sequences in the three-dimensional chromosomal space. After a promoter is identified, RNAP moves through the stages of transcription: open-complex formation, promoter clearance, transcription elongation, termination, and completion of the cycle by returning to promoter search(Borukhov & Nudler, 2008).

The kinetics of transcription has been extensively studied using reconstituted systems *in vitro*, which has yielded mechanistic understandings of the transcription cycle(Henderson et al., 2017; F. Wang et al., 2013; G. Wang et al., 2016). Yet, a living cell is a complex entity; the heterogenous cellular environment is drastically different from the well-mixed solution *in vitro*. Cellular factors such as chromosomal DNA organization(Dame et al., 2020), molecular crowding(Minton, 2001), and regulatory proteins(Seshasayee et al., 2011)may impact the kinetics of promoter search and transcription *in vivo*(Stracy & Kapanidis, 2017). The bacterial nucleoid, composed of chromosomal DNAs, DNA-binding proteins, and RNAs, imposes a unique environment for RNAP to search for promoters and transcribe genes. The nucleoid is neither membranebound nor clearly phase-separated from the cytoplasm(Feric & Misteli, 2021). A recent study finds that the nucleoid behaves like a chromosomal mesh with a pore size of ~ 50 nm, effectively relegating larger complexes to the cytoplasm, such as polysomes that form co-transcriptionally on highly active genes (Xiang et al., 2021). Small, non-DNA-interacting proteins can diffuse through the nucleoid without changes in their mobility. Small, non-specific DNA-interacting proteins, such as the abundant nucleoid associated protein HU, remain nucleoid-localized due to their transitory and weak interactions with the chromosomal DNA (Bettridge et al., 2021; Macvanin & Adhya, 2012; Yamazaki et al., 1984). As the RNAP holoenzyme complex is ~ 20 nm in size (Chen et al., 2017), smaller than the nucleoid mesh pore size, and has significant nonspecific DNA interactions, it likely interacts with the entire volume of the nucleoid during promoter search. The cell metabolic state and transcription profile during different growth conditions and environmental stimuli would influence RNAP’s *in vivo* transcription kinetics as well (Covert & Palsson, 2002; Sastry et al., 2019). Therefore, it is important to study RNAP’s promoter search and transcription kinetics in a realistic cellular environment.

Recent advances in single-molecule live cell imaging technologies have made it possible to track single DNA-binding protein molecules with high temporal resolution and localization precision in small bacterial cells(Bohrer & Xiao, 2020). These emerging studies have generated new insights in DNA-binding proteins’ interactions with the nucleoid and their functions (Bettridge et al., 2021; Geffroy et al., 2022; Stracy & Kapanidis, 2017; Uphoff et al., 2013). For example, studies on the diffusive behavior and spatial localization of RNAP in *E. coli* have identified clustered RNAP organization (Ladouceur et al., 2020; Stracy et al., 2015; Weng et al., 2019) and quantified subpopulations of RNAP molecules that are either freely diffusing, stably binding DNA, or nonspecifically interacting with the nucleoid, each of which would correspond to different stages of promoter search or transcription(Bakshi et al., 2013; Stracy et al., 2015). The spatial distribution of RNAP also changes in response to nutrient rich or poor growth conditions, an effect that cannot be simulated *in vitro* (Ladouceur et al., 2020; Stracy et al., 2015; Weng et al., 2019).These pioneering studies demonstrate the power of singlemolecule imaging in live cells and pave the way to the current study where the kinetics of RNAP during the transcription cycle, in addition to steady-state distributions as in previous studies, is the central focus.

In this study, we probed the kinetics of RNAP’s promoter search and transcription processes under a variety of different conditions in live *E. coli* cells. Using single-molecule tracking (SMT), we confirmed the presence of subpopulations of RNAP molecules engaged in different states of promoter search as previously demonstrated, and further revealed a kinetic path of freely diffusing RNAP molecules routed through rapid, transitory and nonspecific interactions within the nucleoid before binding to DNA more stably. To probe transcription kinetics, we performed fluorescence recovery after photobleaching (FRAP) and determined *in vivo* rates for RNAP’s promoter binding/unbinding, promoter escape, and termination using a kinetic model. Interestingly, we found that transcription kinetics, in particular the promoter escape rate and the percentage of RNAP actively engaged in transcription are sensitive to growth condition, promoter class, and transcription inhibition, but promoter search kinetics are largely independent of these factors. Our work establishes a foundation for further mechanistic studies of bacterial transcription in live cells.

## Results

### Single-molecule tracking (SMT) reveals heterogenous diffusive dynamics of RNAP in live E. coli cells grown in a rich defined medium

To capture the kinetics of RNAP’s search process, we performed single molecule tracking (SMT) using a functional RNAP-PAmCherry fusion strain in which the fusion gene *ropC-PAmCherry* replaces the endogenous chromosomal *ropC* gene encoding for RNAP’s *β*’ subunit. This strain has been previously validated for its functionality in our lab (Weng et al., 2019). The use of the photoactivatable fluorescent protein PAmCherry allows us to activate single RNAP-PAmCherry molecules continuously and stochastically to resolve single molecule trajectories one at a time in single cells (Bettridge et al., 2021; Weng et al., 2019). We grew cells in a rich defined medium (EZRDM) and collected single molecule trajectories with a frame rate of approximately 150 Hz (Δt = 6.74ms). This frame rate enables the tracking of molecules with an apparent diffusion coefficient, *D*_app_, up to 3 μm^2^/s (Supplemental Information). Additionally, the localization uncertainty for a single molecule (~27 nm, **Fig. S1**) in fixed cells set the lower bound of observable diffusion coefficient at *D*_app_ ≥ 0.035 μm^2^/s (Supplemental Information).

We collected a total of 63,182 individual RNAP-PAmCherry trajectories, with an average length of 3.4 ± 2.4 frames from 353 cells (**Fig. S2**). This average trajectory length is shorter than that of the photobleaching curve in fixed cells (6.0 ± 5.9 frames, N =10,727 trajectories from 45 cells, **Fig. S2**), indicating that the tracking trajectories were not limited by photobleaching. RNAP-PAmCherry showed heterogeneous diffusive behavior, with some slow (light blue), some fast (dark blue), and some switched between slow and fast diffusion (yellow) (**Fig. 1A**). Individual trajectories exhibited a wide distribution of *D*app, ranging from 0.1 μm^2^/s to 2 μm^2^/s (**Fig. 1B**), indicating a heterogenous mixture of RNAP sub-populations between which molecules dynamically transition.

**Figure 1:**
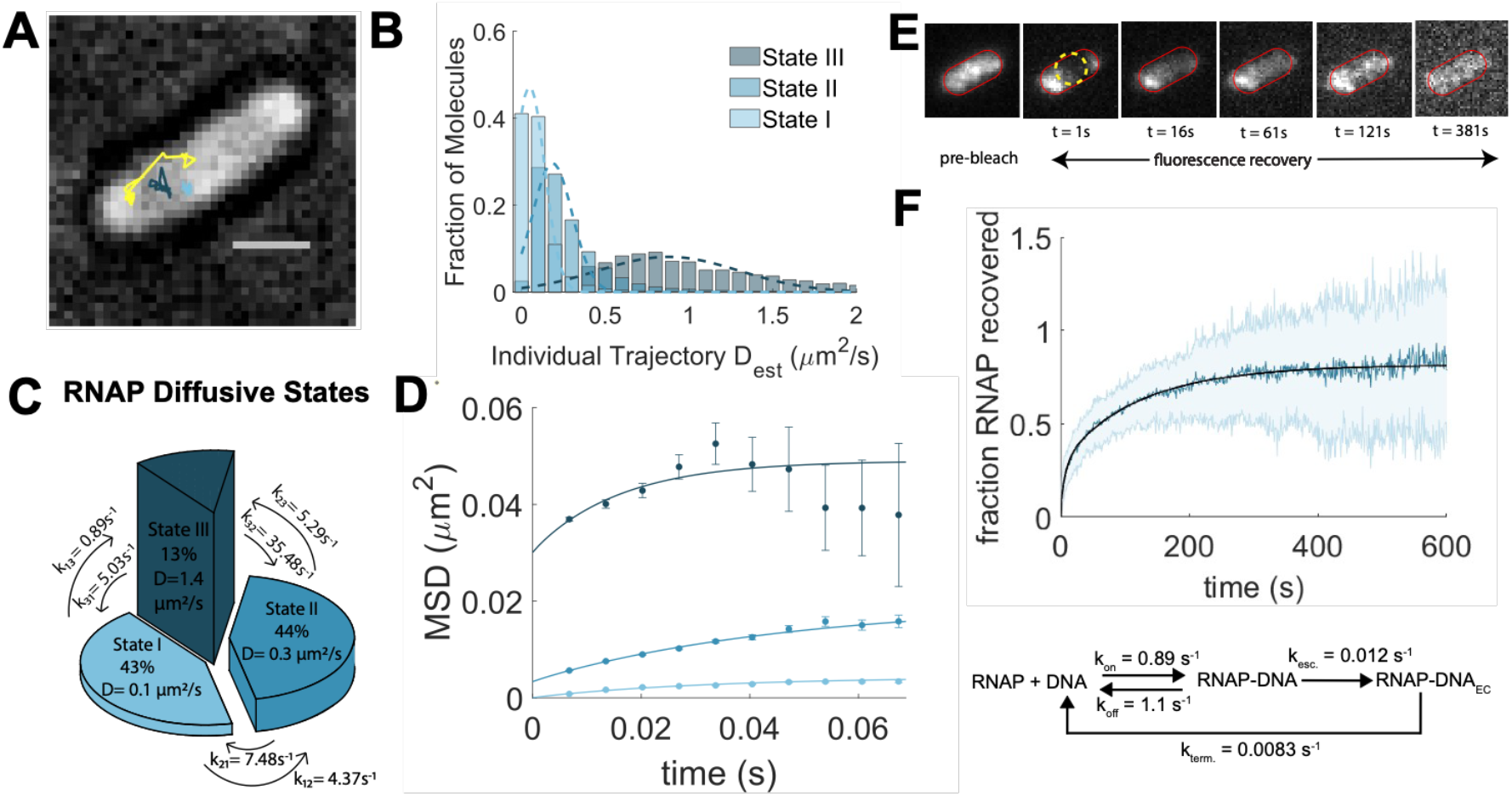
Single-molecule tracking (SMT) to probe RNAP search dynamics and Fluorescence Recovery after Photobleaching (FRAP) to probe RNAP transcription dynamics. A. Representative SMT trajectories of three RNAP molecules overlaid with the corresponding brightfield image of an *E. coli* cell: a slow-diffusing RNAP molecule (light blue), a fast-diffusing RNAP molecule (dark blue), and an RNAP molecule switching between fast-and slow-diffusing states (yellow). Scale bar: 1μm. **B**. Distribution of apparent diffusion coefficients of SMT trajectories. A sum of three Gaussian functions best fit the distribution, corresponding to the three states identified in the Hidden Markov Model (HMM) in **C. C**. Three diffusive states and the corresponding transition rate constants in between identified by the HMM analysis. Pie slice area represents the population fraction, pie slice height the diffusion coefficient, and arrows in between pie slices the transition rate constants (per second per RNAP molecule, **Table 1**). **D**. Mean squared displacement (MSD) curves of SMT trajectories sorted based on the three diffusive states identified by HMM. Color scheme is the same as in **B** and **C**. **E**. A representative FRAP time lapse movie montage with the yellow circle indicating the photobleached area. **F**. Averaged FRAP curve of RNAP-GFP in rich defined media (blue) from 45 cells with the shaded light blue region as standard deviation. The fitted curve(black) from the kinetic model (bottom) with corresponding fit parameters is overlaid on top of the average FRAP curve.

### A Hidden Markov Model analysis identifies three diffusive populations of RNAP

To quantify the dynamic diffusive behaviors of RNAP-PAmCherry molecules, we analyzed SMT trajectories using vbSPT, a variational Bayesian implementation of a hidden Markov model (HMM) (Persson et al., 2013). This method uses thousands of SMT trajectories simultaneously to determine the optimal number of diffusive states, the corresponding diffusion coefficients, and most importantly, the state transition probabilities, from which one can obtain the kinetic rate constants of transitions.

The optimal HMM identified three RNAP diffusive states: state I with *D*_1_ = 0.1 ± 0.002 μm^2^/s, *p*_1_ = 43.5 ± 1.2%; state II with *D*_2_ = 0.3 ± 0.007 μm^2^/s, *p*_2_ = 43.7 ± 1.1%; and state III *D*_3_ = 1.4 ± 0.03 μm^2^/s, *p*_3_ = 12.8 ± 0.4%, μ ± sem, *n* = 63,182 trajectories (**Fig. 1C**). The corresponding dwell times at each state are 198 ± 20 ms, 86 ± 6 ms, and 29 ± 1 ms for States I, II, and III molecules, respectively (**Table S1**). The identification of three diffusive states was consistent with a model-free single-step displacement analysis **(Fig. S3)**. The mean-squared displacement (MSD) of RNAP-PAmCherry molecules in the three states showed clear confined diffusions with progressively larger confinement zones from 161 ± 0.08 nm in size for State I molecules to 313 ± 0.3 nm for State II molecules, and finally to 337 ± 1.1 nm for State II molecules (**Fig 1D**).

The diffusive properties of state I RNAP-PAmCherry matched our expectations for proteins bound to DNA. In our previous work, both a TetR-PAmCherry fusion that binds specifically to the chromosomal DNA and the less diffusive subpopulation of a HU-PAmCherry fusion that specifically binds to DNA structures showed similar confinement zone sizes (~ 200 nm and ~ 230 nm respectively) and apparent diffusion coefficients (*D*_app_ at ~ 0.09 μm^2^/s and 0.14 μm^2^/s respectively) as State I RNAP-PAmCherry molecules (Bettridge et al., 2021). Previous SMT of RNAP molecules also revealed a similar diffusion coefficient (Stracy et al., 2015). The cellular distribution of RNAP-PAmCherry mimics that of HU-PAmCherry across the entire nucleoid, rather than that of TetR-PAmCherry as one concentrated focus (Bettridge et al., 2021), suggesting that the binding sequences of state I RNAP molecules are distributed throughout the nucleoid. The short ~ 200 ms dwell time of State I RNAP molecules (**Table S1**) suggests that the dynamics of this state likely represent some transitory, initial interactions between RNAP with promoters or promotorlike sequences during the search process, but not stable promoter-RNAP complex formation or active transcription, which would be on a much longer time scale base on *in vitro* studies (seconds to tens of seconds (F. Wang et al., 2013)).

By similar comparisons to our previous analysis of the diffusive behaviors of HU-PAmCherry, we reason that State II RNAP-PAmCherry molecules likely represent the subpopulation that randomly diffuses inside the nucleoid with transitory, nonspecific DNA interactions. This subpopulation of RNAP-PAmCherry molecules, with a *D*_2_ ~ 0.3 μm^2^/s, a confinement size of ~ 300 nm (**Fig. 1D**, State II medium blue line), and a short dwell time (~ 90 ms), has nearly identical diffusive behaviors to what we observed on the nonspecific interactions between HU and the nucleoid (*D_app_* = 0.38 μm^2^/s, confinement zone at ~ 300 nm, and a dwell time of ~ 75 ms) (Bettridge et al., 2021). The diffusion coefficient is also similar to what previously observed on RNAP (Stracy et al., 2021) and DNA polymerases (Liao et al., 2016; Uphoff et al., 2013), indicating that these transitory interactions are not specific to DNA sequences but likely a universal dynamics for nonspecific DNA-interacting proteins.

Finally, State III RNAP-PAmCherry molecules had the fastest diffusion coefficient of 1.4 μm^2^/s and the largest confinement zone of ~ 340 nm (**Fig. 1D**, state III dark teal line). Given the size of RNAP (~20 nm, (Chen et al., 2017)) and the radius and diffusion coefficient of free GFP (2.8 nm (Terry et al., 1995) and 7.7μm^2^/s (Elowitz et al., 1999)), the experimentally measured diffusion coefficient of state III RNAP-PAmCherry molecules is close to the predicted value using the Stokes-Einstein equation in a crowded bacterial cytoplasm (~2 μm^2^/s, Supplementary Information), suggesting that this population of RNAP molecules diffuses freely in the cytoplasmic of the cell. Consistent with this suggestion, a previous study found that in the DNA-free endcaps of replication-stalled cells, RNAP molecules had a mean diffusion coefficient of ~ 2 μm^2^/s (Stracy et al., 2015).

### Transition analysis identifies a preferred kinetic flow for promoter search

A major advantage of the HMM analysis of trajectories over other methods, such as a Gaussian mixture model of trajectory *D*_app_ or single step displacement analysis, is the calculation of state transition probabilities and the resulting state transition rate constants (Persson et al., 2013). As shown in **Fig. 1C**, the kinetic rate constants among the three states ranged from ~ 1 s^-1^ to ~ 35 s^-1^ (**Table S1**). These rapid transition kinetics indicate that RNAP molecules detected under our imaging conditions are interacting with DNA and the cell milieu transiently, consistent with the expectation of promoter search.

Interestingly, the analysis revealed a preferred kinetic path for freely diffusing RNAP molecules to bind to DNA. Freely diffusing RNAP molecules (State III) enter state II faster than state I (compare *k*_32_ at 35.5 ± 2.4 s^-1^ and *k*_31_ at 5.0 ± 1.2 s^-1^), suggesting that freely diffusing RNAP rarely bind DNA directly. After entering the nucleoid and interacting with the nucleoid nonspecifically (state II), RNAP transitions into the DNA-bound state I with a larger rate constant (*k*_21_ = 7.5 ± 0.8 s^-1^) than that of returning to the freely diffusing state III (*k*_23_ = 5.3 ± 0.6 s^-1^). DNA-bound RNAP molecules (state I) are more likely to return to the nucleoid-diffusing state II (*k*_12_ = 4.4 ± 0.6 s^-1^) than the freely diffusing state III (*k*_13_ = 0.9 ± 0.2 s^-1^). As such, the majority of RNAP molecules (~ 90%, state I plus state II) are retained in the nucleoid for continuous promoter search. Assuming that RNAP makes contacts with all chromosomal sequences in the E. coli genome during promoter search, we calculated the apparent on-rate of RNAP to nonspecific DNA at ~4 x 10^5^ M^-1^s^-1^, and the apparent *K_d_* at 2mM for nonspecific DNA sequences (Supplemental Information, **Table S1**). These results suggest that RNAP prefers to stay nucleoid localized and that there is a preferred kinetic path from freely diffusing RNAP to the DNA-bound state, implicating that the RNAP search strategy is limited by non-specific DNA interactions.

### FRAP of RNAP reveals transcription kinetics in live cells

SMT experiments described above allowed us to access the rapid promoter search kinetics with a high temporal resolution. While some DNA-bound, state I RNAP molecules could be transcribing, the millisecond time scale of SMT does not allow the probing of transcription kinetics, which ranges from tens of seconds to minutes. Therefore, we employed FRAP (fluorescence recovery after photobleaching) to evaluate RNAP’s *in vivo* transcription kinetics.

We grew MG1655 *E. coli* cells expressing a GFPuv labeled RNAP (fused to the β’ subunit(Cabrera & Jin, 2003)) under the same rich defined medium (EZRDM) growth condition as that in SMT. We photobleached a diffraction-limited area in a cell’s nucleoid and quantified the subsequent fluorescence signal recovery rate over a period of 10 min with a frame rate of 1 Hz. At this time scale and sampling rate, the fluorescence signal recovery, caused by the entry of fluorescent RNAP-GFP molecules and exit of photobleached RNAP-GFP molecules in this area, is due to the transcription cycle (initiation, elongation and termination cycle) in addition to the random diffusion process (**Fig. S4**).

Using a kinetic scheme that reflects both the random diffusion and the transcription cycle (**Fig. 1F**), we determined best-fit parameters through least squares minimization of simulated FRAP curve to the experimentally measured FRAP curve. We found that on average, 34% of RNAP-GFP randomly diffused, 27% bound to DNA, and 40% actively engaged in transcription within the photobleached area (**Figure 1E**, **Table S3**). The random diffusion population contributes significantly to the rapid recovery of the fluorescence within the first few seconds of the FRAP curve.

Most importantly, we were able to extract transcription kinetic parameters from the FRAP curve using the model. As shown in **Fig. 1F**, RNAP binds and unbinds DNA with an apparent *k_on_* and *k_off_* at ~ 1 s^-1^. Because these rate constants are ~ 5 times slower than the transitory DNA-binding/unbinding kinetics between state I and II RNAP molecules observed in SMT (**Fig. 1C**), we reason that they most likely reflect the formation of the RNAP-DNA closed complex, where RNAP can dissociate to initiate another round of promoter search and binding, or proceed to the formation of open complex, which is a prerequisite for successful transcription initiation. The subsequent rate constant *k_esc_*, which we tentatively assign as the promoter escape rate, is at 0.012 s^-1^, similar to the promoter clearance transition rate previously measured in biochemical studies(Fazal et al., 2015; Vo et al., 2003). This rate constant suggests that it takes on average about 1/0.012 ≈ 80 s for a promoter-bound RNAP molecule to clear the promoter to enter the elongation phase. The last rate constant, *k*_term_ is = at 0.0083 s^-1^, corresponds to an average dwell time of ~120 seconds, nearly identical to the previously measured average rRNA operon transcription time of ~130 seconds(Gotta et al., 1991), suggesting that this step is likely the average transcription elongation time of active operons. Note that a simpler kinetic model that only describes the binding and unbinding of RNAP to DNA without involving a transcription elongation process could not describe the observed recovery dynamics at the much slower time scale (**Fig. S7**).

### An RNAP mutant defective in open-complex formation exhibits only promoter search dynamics

Characterizations of the promoter search and transcription kinetics of RNAP under the wildtype (WT) condition allowed us to establish a baseline to further explore factors contributing to the observed kinetics. We reason that if the dynamics of RNAP observed in SMT indeed reflects the promoter search process but not transcription kinetics, we should observe a similar behavior on an RNAP mutant that does not proceed to transcription. To examine this possibility, we constructed a PAmCherry fusion to a non-transcribing mutant of RNAP, RNAP^I1309A^. The RNAP^I309A^ mutant has been shown *in vitro* to retain the ability to form a holoenzyme with σ^70^ but has significantly impaired ability to form open, transcribing complexes,^10^ limiting the DNA-RNAP interaction to DNA binding and unbinding without transcription. Using co-immunoprecipitation, we confirmed that the RNAP^I309A^-PAmCherry fusion is incorporated into the holoenzyme when induced from a plasmid (**Fig. S5**).

SMT and subsequent HMM analysis showed that the RNAP^I1309A^ mutant has largely similar diffusive behaviors as WT RNAP-PAmCherry. State I RNAP^I309A^-PAmCherry has a diffusion coefficient *D*_1_ at 0.1 ± 0.003 μm^2^/s (50 ± 2%) and State II *D*_2_ at 0.38 ± 0.007 μm^2^/s (50 ± 2%, N =13,473 trajectories from 275 cells, **Fig. 2A**). The corresponding confinement zones of the two states and dwell times are slightly larger than those for WT (**Fig. 2B-E, Table S2**), likely due to the enhanced non-productive interactions between this RNAP mutant with DNA (Zhang et al., 2015). We did not observe the minor state III, the fastest diffusing subpopulation of WT, in our RNAP^I309A^-PAmCherry data. It is possible that state III molecules may not be statistically well represented by the much smaller sample size of trajectories collected on this mutant and hence likely mixed with state II molecules. Consistent with this possibility, the transition rate constant of RNAP^I309A^-PAmCherry from state II to state I (*k*_21_ = 19.5 s^-1^) is larger than that of the WT (*k*_21_ = 7.5 s^-1^), likely resulting from the combined process that integrates both the entry of freely diffusing RNAP molecules (state III) into the nucleoid interacting state (state II) and the subsequent DNA-binding (state I). Nevertheless, the largely similar diffusive behaviors between the RNAP^I309A^ and WT RNAP suggest that they primarily reflect RNAP search kinetics but not transcription processes.

**Figure 2.**
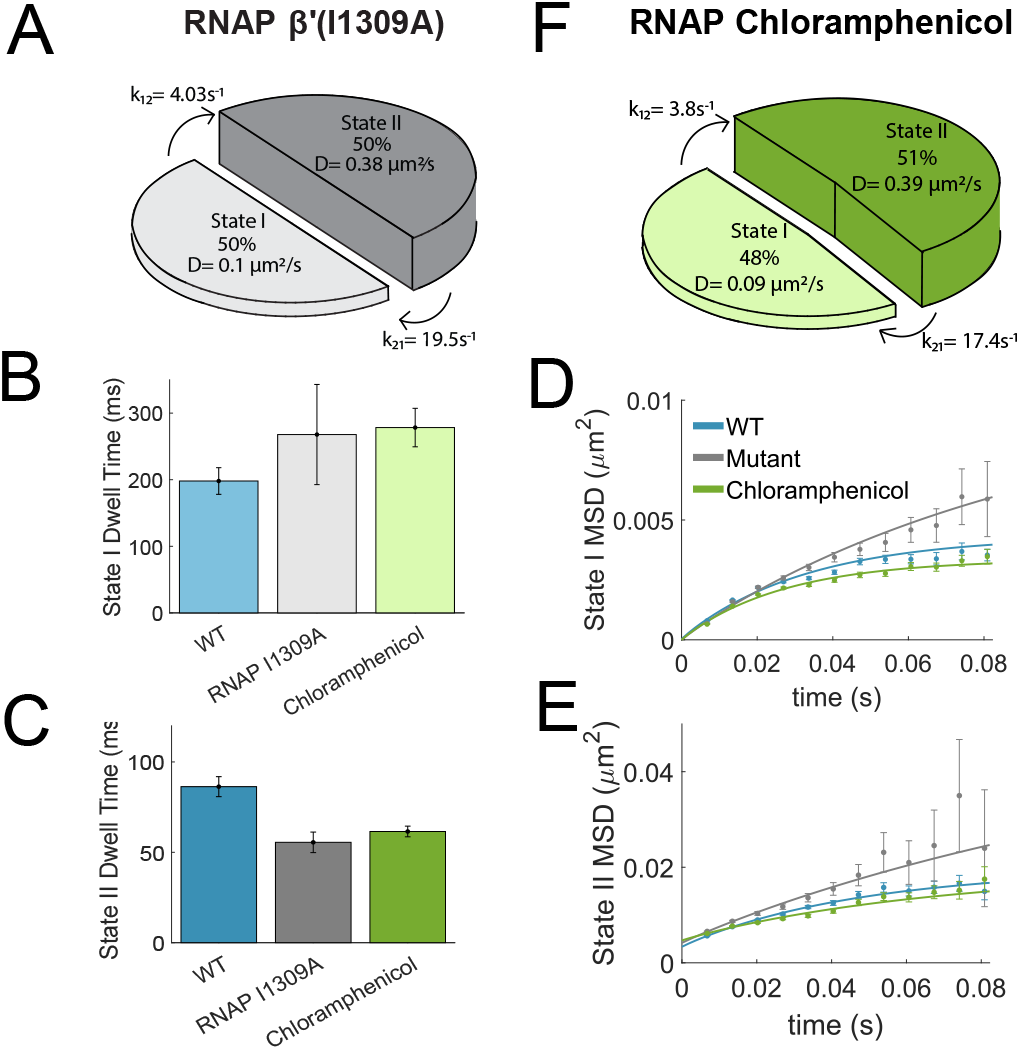
RNAP promoter search kinetics is not affected by an RNAP mutant that has impaired transcription initiation (A) or chloramphenicol treatment that leads to nucleoid compaction (B). **A**. Pie chart representation of the SMT result for RNAP β’(I1309A). **B** and **C**. Average dwell time for state I (**B**) and state II (**C**) RNAP molecules for RNAP β’(I1309A, gray) and chloramphenicol-treated (green) cells in comparison with WT cells (cyan). **D** and **E**. MSD plots for state I (**D**) and state II (**E**) RNAP molecules for RNAP β’(I1309A, gray) and chloramphenicol-treated (green) cells in comparison with WT cells (cyan). Error bars represent s.e.m. **F**. Pie chart representation of the SMT result for RNAP in cells treated with chloramphenicol (600 μg/ml, 30 min).

### RNAP search kinetics are not affected by nucleoid compaction

Next, we examined the effect of nucleoid density on the promoter search kinetics of RNAP. In chloramphenicol treated cells, the nucleoid compacts significantly(Bakshi et al., 2012, 2014; Cabrera et al., 2009; van Helvoort et al., 1996; Zimmerman, 2006), which lead to reduced nucleoid mesh pore size(Xiang et al., 2021). However, most transcription, including ribosome RNA operons(Dennis, 1976), continues except for a few transertion-dependent operons such as the *lac* operon(Graham et al., 1982). Therefore, we can isolate the effect of nucleoid compaction from transcription. We treated cells with chloramphenicol at a concentration of 600 μg/ml for 30 minutes and observed clear nucleoid compaction (**Fig. S6**) performed SMT on RNAP-PAmCherry showed similar diffusion dynamics of RNAP-PAmCherry under this condition compared to that of WT and the RNAP^I130A^-PAmCherry mutant (**Fig. 2F**, *D*_1_ = 0.09 ± 0.001 μm^2^/s, 48.1 ± 0.7% for state I molecules, *D*_2_ = 0.39 ± 0.003 μm^2^/s, 51.9 ± 0.7%, N = 44,168 trajectories from 121 cells). This result suggests that the nucleoid diffusion and binding dynamics of RNAP did not change significantly when the nucleoid is compacted.

Interestingly, the dwell time of state I RNAP-PAmCherry molecules increased from ~ 200 ms in WT to ~280 ms with chloramphenicol-induced compaction and remained similar to that of the RNAP^I309A^-PAmCherry mutant (**Fig. 2B**). As the dwell time of state I molecules does not depend on the concentration of RNAP but reflects the time it takes for a bound RNAP molecule to dissociate from DNA, the increased dwell time suggest that under the condensed nucleoid condition, local high DNA density may prevent rapid dissociation of DNA-bound RNAP molecules, or that there exist multiple consecutive, repetitive rebinding events that are undistinguishable at the time resolution of SMT. Previous theoretical biophysical models of the search process of DNA-binding proteins demonstrate that crowding can indeed influence the dissociation rate of proteins bound to the DNA(Bauer & Metzler, 2012; Zaid et al., 2009).

### Poor growth medium affect transcription initiation kinetics but less on promoter search or elongation kinetics

Previous studies have documented that translation and transcription profiles are impacted by cell growth in media of different nutrient levels(Bhatia et al., 2022; Li et al., 2014). Mechanistically, a recent study shows that the translational elongation rate by ribosome scales linearly with the growth rate through the sensing of ppGpp level in cells(Wu et al., 2022). How transcription kinetics responds to different growth rates is unknown.

To investigate this question, we performed SMT of RNAP-PAmCherry and FRAP of RNAP-GFPuv in cells grown in a minimal defined medium M9 supplemented with glycerol to compare with experiments described above, where cells were grown in a rich defined medium (EZRDM). We observed that RNAP-PAmCherry exhibited two similar diffusive states (*D*_1_ at 0.11 ± 0.003 μm^2^/s, *p*_1_ = 44.2 ±1.3% for state I and *D*_2_ at 0.38 ± 0.005 μm^2^/s, *p*_2_ = 55.8 ±1.3, N =21,500 trajectories from 127 cells) with similar dwell time and diffusive domain sizes, but a faster transition rate from nucleoid interacting state II to DNA-bound state I (*k*_21_ = 11.6 ± 1.2 s^-1^, **Fig. 3A, Fig S7, Table S1**). The FRAP data showed a larger fraction of freely diffusing RNAP molecules (41.9%) and a smaller fraction of promoter-bound, non-transcribing RNAP (18.7%, N = 46 cells, **Fig. 3B**) compared to the rich growth condition, suggesting a reduced transcription activity at the nutrient-limited state. Accordingly, the transcription kinetic model of the FRAP curve (**Fig. 3B**) revealed that the promoter-binding rate was reduced and the unbinding rate enhanced compared to what were observed in EZRDM medium (*k_on_* = 0.63 s^-1^ and *k_off_*=1.4 s^-1^ respectively compared to *kon* = 0.89 s^-1^ and *k_off_* =1.1 s^-1^ under EZRDM rich growth condition). Since the SMT data did not differ from that of WT, the difference in the apparent *k_on_* and *k_off_* is likely caused by the reduced number of available promoters, and/or that the failure of promoter-bound RNAPs to initiate successfully. Interestingly, the apparent promoter escape rate *k_esc_* (0.019 s^-1^ or ~ 50 s to clear a promoter) is about 60% faster than that in EZRDM (*k_int_* = 0.012 s^-1^ or ~ 80 s to clear a promoter), while the apparent termination rate *k_term_* (0.0092s^-1^, corresponding to ~ 110 s average elongation time) is only marginally slower than that in EZRDM (*k_term_* = 0.0083 s^-1^ or ~ 120 s). These results suggest that the nutrient-limited growth condition does not affect promoter search or transcription elongation, but the formation of productive promoter-RNAP complexes and the subsequent promoter escape step.

**Figure 3.**
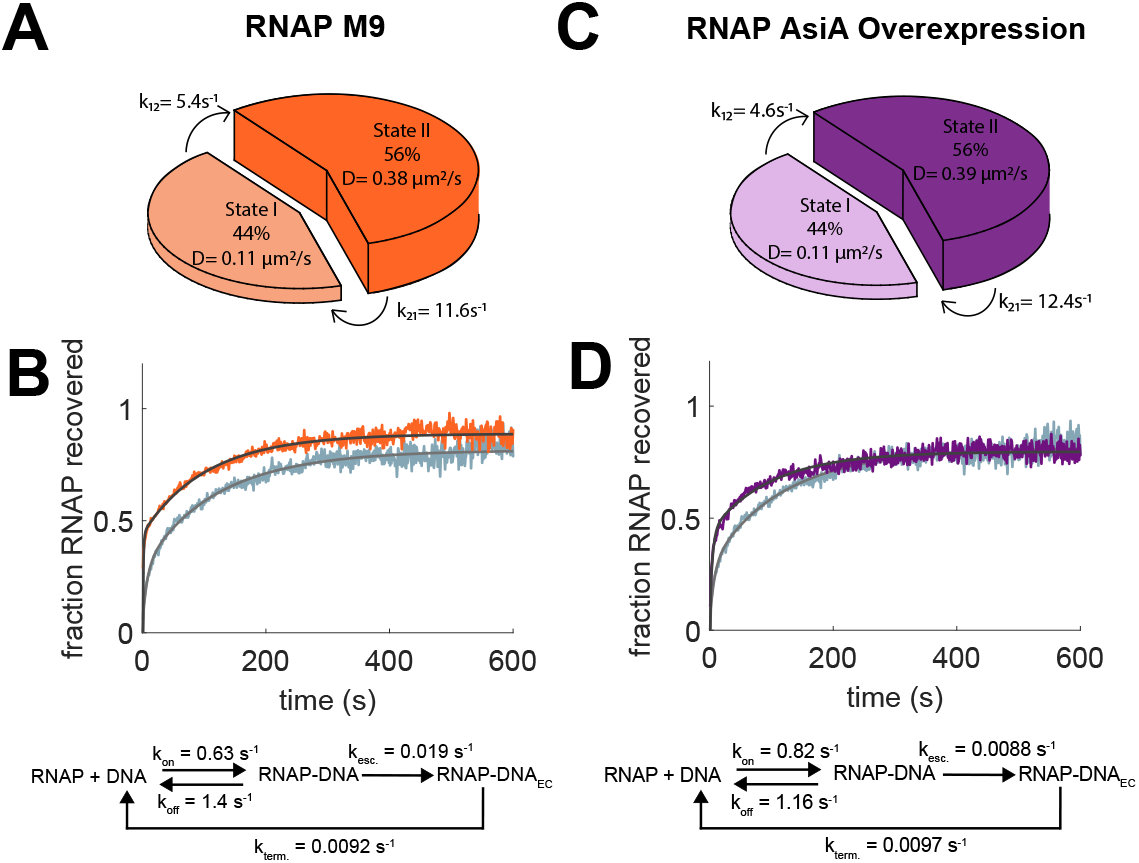
RNAP transcription kinetics, but not promoter search kinetics, is sensitive to growth condition and the use of alternative promoters. **A** and **C**. Pie chart representations of the SMT results for cells grown in M9 minimal media (**A**) and cells overexpressing AsiA (**C**). **B** and **D**. Averaged FRAP curves of RNAP-GFP in M9 minimal medium from 46 cells (**B**) and in cells overexpressing AsiA from 45 cells (**D**). The fitted curves (black) from the kinetic models (bottom) with corresponding fit parameters is overlaid on top of the average FRAP curve. The FRAP curve of WT cells in EZRDM rich medium is shown in light cyan as a comparison.

### RNAP exhibits slowed promoter escape rate on alternative promoters

Under the normal exponential growth conditions, σ^70^ accounts for 65-95% of the total cellular pool of σ factors, mainly responsible for the transcription of ribosomal operons and housekeeping genes containing classic −10/−35 promoters(Jishage et al., 1996). To investigate the promoter search and transcription kinetics of alternative promoters and σ factors, we overexpressed the T4 phage anti-σ^70^ protein AsiA(Orsini et al., 1993). AsiA binds to σ^70^ tightly and form a ternary complex with core RNAP to redirect the modified holoenzyme from recognizing classic 10/-35 promoters to phage promoters(Hinton et al., 2005; Severinova et al., 1998; Sharma & Chatterji, 2008). As such, in the absence of T4 phage infection, the modified Asia-RNAP holoenzyme is forced to use alternative promoters, many of which are −10 class promoters (containing extended −10 sequence and lacking −35 sequence)(Severinova et al., 1998). Unbound core RNAP may also complex with alternative σ factors to direct expression of other subset of genes.

We performed SMT of RNAP-PAmCherry and FRAP of RNAP-GFPuv in cells overexpressed AsiA from a plasmid (0.2% arabinose induction for 2 hours). The promoter search kinetics under this condition resembled that in the M9 medium in nearly all aspects (**Fig. 3C, Fig. S7, Table S1**), suggesting that the search kinetics of RNAP is independent of promoter sequences and the identity of alternative σ factors. However, the FRAP curve deviated from that in M9 and EZRDM media (**Fig. 3D**). The percentage of randomly diffusing RNAP molecules (43%, N = 45 cells) was similar to that in M9 medium and higher than that in EZRDM, but percentage of RNAP molecules actively engaged in transcription elongation phase (27%) appeared to be lower than both M9 and EZRDM conditions (**Table S3**). Most interestingly, the promoter binding/unbinding and elongation kinetics of RNAP in the presence of AsiA overexpression are similar to those in the EZRDM condition, but the apparent promoter escape rate *kesc* (0.0088s^-1^, or ~ 115 s to clear a promoter) is reduced ~ 30% compared to that in EZRDM. These results suggest that under our experimental conditions, transcription is mostly modulated at the level of promoter escape and the percentage of RNAP engaged in active transcription, while promoter search and transcription elongation kinetics remain independent of growth conditions and genes.

### RNAP search and transcription kinetics are influenced by rifampicin

Rifampicin is a well-established transcription inhibitor. It binds to the RNA exit channel of the β-subunit, blocking the exit of the transcript when it grows to 2-3 nucleotides (nts) (Campbell et al., 2001). As such, RNAP is unable to escape the abortive initiation phase and remains trapped on the promoter(Herring et al., 2005). Additionally, rifampicin inhibition leads to the expansion of the nucleoid due to the rapid degradation of cellular mRNAs(Bakshi et al., 2014; Cabrera et al., 2009; Cabrera & Jin, 2003; Stracy et al., 2015). Therefore, we reason that this condition offers another window to examine how the promoter search and transcription kinetics of RNAP are impacted due to the occlusion of promoters from diffusing RNAP molecules and the change of the nucleoid environment.

We performed SMT of RNAP-PAmCherry and FRAP of RNAP-GFPuv in cells treated with 200 μg/mL rifampicin for 15 minutes. We observed clear nucleoid expansion in these cells (**Fig. S8A**), indicating the fast effect of rifampicin. Interestingly, we observed that both state I and II RNAP molecules in rifampicin-treated cells become more diffusive, with *D*_1_ and *D*_2_ increasing from ~0.1 to ~ 0.2 μm^2^/s (0.17 ± 0.004 μm^2^/s) and from 0.3 μm^2^/s to ~ 0.8 μm^2^/s (0.83 ± 0.008 μm2/s, N = 40,144 trajectories from 361 cells) respectively (**Fig. 4A**). Correspondingly, the diffusive domain increased for each state (**Table S2**). While the dwell time for state I decreased, there was no significant change in the dwell time of state II (**Fig. S8B, Table S1**). These results suggest that RNAP becomes much more diffusive in the nucleoid, either because of the expanded nucleoid volume and/or that there are very few unoccupied promoter sequences available for RNAP to make initial nonspecific contacts.

**Figure 4:**
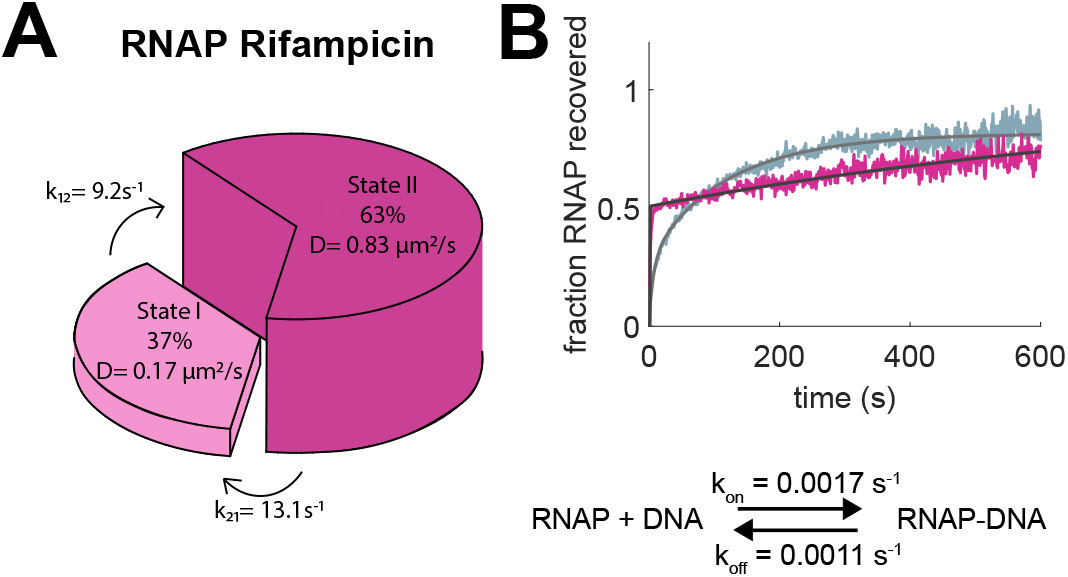
Rifampicin changes both promoter search and transcription kinetics by altering RNAP’s nonspecific and specific DNA interactions. **A**. Pie chart representation of SMT result of RNAP in cells treated with rifampicin (15 μg/ml, 15 min). **B**. Averaged FRAP curve of RNAP-GFP for cells treated with rifampicin (pink, n=46). The fitted curve (black) from the kinetic model (bottom) with corresponding fit parameters is overlaid on top of the average FRAP curve. The FRAP curve of WT cells in EZRDM rich medium is shown in light cyan as a comparison.

Most interestingly, the FRAP curve exhibited a rapid recovery phase (~2 s) as we observed in other conditions, followed by a much slower recovery phase that only reached about 75% of the initial intensity after 500 s (**Fig. 4B**). The transcription kinetic models revealed that there are 39% of RNAP molecules freely diffusing and 61% of RNAP molecules trapped on promoters, of which ~ 25% of them do not recover over the 10 minutes of the FRAP experiment (N = 45 cells). The transcription kinetic model of the FRAP curves showed a one-step binding and unbinding reaction with DNA, with an apparent *k_on_* at 0.0017s^-1^ and *k_off_* at 0.0011 s^-1^, consistent with RNAP slowly exchanging on the DNA (~ 1 RNAP molecule trapped every ~ 600 s and dissociates every ~ 900 s) (**Fig 4B**). The significantly divergent fluorescence recovery and kinetic model outcomes of rifampicin-treated cells demonstrates that rifampicin diminishes the transcription elongation phase as was observed under the WT condition but preserves the initial promotor binding step.

## Discussion

In this work, we investigated the promoter search and transcription dynamics of RNAP in live *E. coli* cells under various conditions. Using single-molecule tracking, we not only confirmed the presence of different diffusive states of RNAP reflecting different species in the process of promoter search as previously demonstrated(Bakshi et al., 2013; Stracy et al., 2015), but also revealed a kinetic path of the search process (**Fig. 1C**). A freely diffusing RNAP molecule first engages in rapid, transitory interactions nonspecifically with the nucleoid in a local domain before it converts to a more stably DNA-bound state, from which it can either proceed to promoter complex formation for transcription or dissociate to diffuse away to initiate another round of promoter search. Rarely, a free diffusing RNAP molecule binds to DNA directly without going through the nonspecific interaction state, indicating the role of nonspecific interactions in facilitating the search process. This flux suggests that an RNAP molecule likely repetitively hops on adjacent DNA segments in a local domain until it encounters a promoter or promoter-like sequence or diffuses out of the domain to initiate another round of repetitive hoping and testing in the nucleoid. This search process is consistent with what has been previously observed on the binding of transcription factors to nonspecific DNA sequence(Afek et al., 2014) and the nucleoid-localization of RNAP(Stracy et al., 2015), suggesting an important role of nonspecific interactions in facilitating sequence-specific binding of RNAP (Stracy et al., 2021). Interestingly, this search process is largely independent of promoter open complex formation (**Fig. 2A**), nucleoid compaction (**Fig. 2B**), growth condition (**Fig. 3A**), or the promoter class in use (**Fig. 3C**), indicating an intrinsic (likely electrostatic), non-sequence-dependent interactions between RNAP and the chromosomal DNAs.

Using FRAP, which allowed us to access the diffusive dynamics of RNAP at a much longer time scale than probed by SMT, we were able to capture the overall kinetics of the transcription cycle in live *E. coli* cells (**Fig. 1C, D**). The transcription rates recovered from the FRAP kinetic model generally agree with previous biochemical measurements. For example, the average *in vivo* transcription rate at 30 °C was found to be ~25 nt/s(Taniguchi et al., 2010). With an assumed slower, room temperature (~24°C) elongation rate of ~15-10 nt/s at our experimental conditions, the average transcription time for a single operon with an average size of ~1000 nt would be ~70-100 seconds, comparable to what were observed in our model. Additionally, unlike RNAP promoter search kinetics, RNAP transcription kinetics are sensitive to altered growth conditions (**Fig. 3B**) and the use of different promoter classes under the AsiA overexpression condition (**Fig. 3D**), primarily responding to altered conditions by changes in the percentage of RNAP actively engaged in transcription and the apparent promoter escape rate (**Fig. 1F, 3B, D**), which are both related to specific promoter sequences.

These distinct results suggest the promoter search process of RNAP is independent of the subsequent transcription cycle, in that nonspecific DNA interactions dominant and facilitate the search process, while specific promoter DNA interactions determine whether and how fast transcription proceeds. The transcription elongation rate appears to be relatively constant under the experimental conditions we have tested, indicating that it may not be a major regulatory step of transcription.

One interesting phenomenon we observed in SMT of RNAP is that the third diffusive state under the rich medium growth condition (EZRDM), or the most diffusive state (*D*_3_ = 1.3 μm^2^/s), did not appear under any other conditions. As this third state constitute only a minor population of total RNAP molecules (p = 12.8 ± 0.4%), It is possible that the largest sample size under the EZRDM condition (63,182 trajectories compared to the next largest sample of 44,168 trajectories under the chloramphenicol condition) made the third diffusive state statistically more prominent than that in the other conditions. This minor population is unlikely caused a higher expression level of RNAP under the rich medium growth condition that exceeds the binding capacity of the nucleoid, as Western blots confirmed that this condition did not have significantly higher expression of RNAP compared to that grown in complex rich medium LB (LB had 1.2 times more expression compared to EZRDM (**Fig. S9**). Additionally, the expression levels of RNAP decreased in M9 to 48% of that of the WT EZRDM condition but increased to about 2-fold under the AsiA overexpression condition (**Fig. S9**), suggesting that RNAP expression level is unlikely a contributing factor for the presence or absence of state III RNAP molecules. An alternative is that this minor population of RNAP could represent a minor RNAP species that has little interactions with DNA, either due to the absence of a σ factor, or the large cytoplasmic volume under the rich medium growth condition. RNAP molecules with a similar apparent diffusion coefficient under a similar rich growth medium condition in *E. coli* cells has indeed been previously observed (*D* = 1.1 μm^2^/s) and identified as non-DNA interacting (Stracy et al., 2015).

Finally, the rifampicin-treated condition represents an interesting deviation from the kinetic scheme described above. We showed that under this condition, ~ 60% of RNAP molecules become more diffusive, and ~ 40% are stably bound. The more diffusive RNAP molecules appeared to have different nonspecific interactions with the nucleoid, as the apparent state II diffusion coefficients molecules are larger than those under other conditions. Perhaps most importantly, the transition rate from the DNA-bound state I to nucleoid diffusing state II (*k_21_* at ~ 9 s^-1^) is about twice faster than those under all other conditions and corresponds to a shortened dwell time of state I molecules (τ = ~ 120 ms in comparison to ~ 200 −300 ms under other conditions). These results suggest a weakened interaction of RNAP with nonspecific DNAs, which could be due to the expanded nucleoid volume where low local DNA densities reduce potential interactions. The 40% stably bound RNAP molecules are presumably trapped on promoters due to the blockage of the RNA exit channel by rifampicin after the synthesis of 2-3 nts. The apparent on-rate (0.0017 s^-1^) of RNAP to the DNA-bound state under this condition is ~ 500 times slower than that of WT, and ~ 7-fold slower than the apparent promoter escape rate under the WT condition, suggesting a significant kinetic barrier for a RNAP-promoter complex to proceed to transcription initiation. The slow off-rate (0.0011 s-1) is also ~ 10-fold slower than the termination rates we observed under other conditions, indicating that the dissociation of rifampicin-trapped RNAP may require mechanisms involved in resolving stalled transcription elongation complexes(Hatoum & Roberts, 2008).

## Conclusions

In this study, we probed quantitatively the *in vivo* RNAP promoter search and transcription kinetics using SMT and FRAP data respectively. We found that RNAP’s promoter search is facilitated by nonspecific DNA interactions and largely independent of nucleoid organization, growth condition, transcription, or promoter classes. RNAP’s transcription kinetics, however, is sensitive to these conditions and mainly modulated at the levels of actively engaged RNAP and the promoter escape rate. Our work establishes a foundation for further mechanistic studies of bacterial transcription in live cells.

## Materials and Methods

### FRAP data collection

For FRAP experiments, cells were inoculated from freshly streaked LB plates into 2mL of EZRDM or M9 media and grown overnight (~16 hours) at 24°C with shaking. Saturated cultures were diluted 1 to 200 into fresh 2mL of their respective media and let grown for several hours to reach mid-log phase growth (O.D.600 ~ 0.4). For rifampicin treatment, rifampicin stock at 50mg/mL was added such that the final concentration was 200 μg/mL and incubated with shaking at 24°C for 15 minutes. For AsiA overexpression, the strain containing the pDR-pBAD-AsiA plasmid was spun down at OD600 ~ 0.25 and resuspended in EZRDM media with glycerol as the main carbon source with 0.2% arabinose added to induce AsiA expression. Cells were induced for 2 hours at 24°C with shaking. Cells in all conditions were washed once then spun and resuspended in 1/20^th^ volume to concentrate the cells.

Harvested cells were pipetted onto agarose pads and sandwiched between the agarose pad and a #1 coverslip as previously described (41). Immobilized cells were imaged on an Olympus IX71 inverted microscope with a 100x oil objective (NA =1.45). Photons from cells were collected with an Andor EMCCD camera using MetaMorph imaging software (Molecular Devices). Fluorescence from cells was obtained using solid-state lasers at 488nm wavelength (Coherent). Twenty frames at 100ms exposure per frame were taken before cells were bleached. Cells were bleached along the quarter cell by focusing the laser light onto the imaging plane. The ratio of the confocal beam and epifluoresence beam intensities was 100 to 1. Usually, an area encompassing ~20% of the total cell area was bleached using this set-up. Fluorescence recovery was monitored by acquiring 100ms fluorescence images every second for 600 frames (10 minutes).

### FRAP data analysis

To account for photobleaching, fluorescence recovery for each cell was calculated by dividing the fluorescence within the bleached region and the cell. To account for heterogeneity of fluorescence within individual cells (Fig. SX), this cell normalized ratio after photobleaching was divided by the same ratio before bleach (i.e. the “steady state” distribution, which was the average distribution over 20 images taken immediately before the bleaching step). The bleached region centroid was determined by the position of the maximum pixel value in the bleach frame, and the region radius was determined to be the full-width half maximum of the confocal spot intensity, ~3 pixels. This radius was used throughout the analysis. The cell region was determined by first rotating the cell such that the long axis corresponded to the y-axis, then selected manually as a rectangular region. Due to incomplete photobleaching, the above normalized fluorescence recovery ratio did not start at zero. To normalize each recovery to start at zero, we first estimated the zeroth timepoint by estimating the derivative at the first timepoint and subtracted this value from the normalized recovery ratio. To normalize such that the recovery could reach the theoretical maximum of one, the normalized recovery ratio was then divided by (1 – min_val) to rescale. Due to heterogeneity between cells, FRAP recoveries were averaged over all cells for each condition.

### FRAP modeling

For FRAP simulations, differential equations describing changes in concentrations of the various states based on the kinetic rates described in the main text were solved numerically using the forward finite difference method for the bleached and fluorescent populations. Freely diffusing molecules were assumed to be distributed equally between the simulated bleached region and the remainder of the cell. Residuals between the simulated FRAP recovery and the averaged FRAP recovery for each condition were minimized using the LSQNONLIN function in MATLAB version 2017a. Minimum values of 0 were imposed for all parameters. Initial guesses were randomized to ensure solutions were not due to convergence on a local minimum. For the EZRDM and AsiA overexpression conditions, two sets of parameters were found to be able to describe the data. For each condition, the parameter set that had the lowest squared residuals was chosen as the final model.

### SMT data collection

Cells were grown and harvested as previously described. For the RNAP fluorescent fusion, the C-terminus of the endogenous β’ subunit of RNAP, RopC, was tagged with the photoactivatable fluorescent protein PAmCherry. The fusion protein was expressed at the expected full-length and supported WT-like cell growth rate as the sole copy of cellular β’ subunit of RNAP (Figure S1). Cells were grown in defined rich medium (EZRDM) at room temperature (RT). For the RpoC(I1309A) mutant, the MG1655 strain containing the pCH-rpoC(I1309A)-PAmCherry plasmid was induced with 1mM IPTG for 2 hours, then resuspended in media without IPTG and let grown at 30°C for one hour to allow for maturation of the PAmCherry protein before harvest as described above. For chloramphenicol treatment, chloramphenicol was added to a final concentration of 600ug/ml for 30 minutes before cells were harvested for imaging. Rifampicin treatment, AsiA overexpression and M9 growth conditions were the same as described for FRAP experiments. Cells were illuminated with solid-state lasers at 405nm and 568nm wavelengths (Coherent). All SMT images were collected using 5ms exposure with 1.74ms of cycle time for a total frame length of 6.74ms. Three movies consisting of 2500 frames each were taken for each cell. To maximize data collection, the UV power was modulated between consecutive movies to maintain a similar activation rate across the imaging acquisition. No cell was imaged longer than five minutes to avoid phototoxicity effects.

### SMT data analysis

For single molecule tracking analysis, tiff stacks of cell images were imported into the single molecule tracking software UTrack version 3.1 (42) within the Matlab 2017a software. We performed the detection and tracking of molecules on individual movies using the Gaussian Mixture Model setting within the UTrack software. For detection, an α value of 0.01 was used; for frame-to-frame linking, 15 frames was the maximum time and 0 to 2 pixels was the maximum distance to link trajectories together. Linked trajectories were then filtered by their intensity (between 0.5-1.5x the average single molecule intensity) and their localization within cells (i.e. any trajectory outside of cells was excluded) and exported as MATLAB files. Trajectories were analyzed using custom inhouse code (available upon request) to assess their diffusive properties. We verified the HMM properties by calculating the cumulative displacement probability and fitting the distribution to this function:

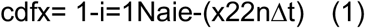

where *x* is the single frame displacement, *N* is the total number of diffusive states, *ai* is the relative fraction of molecules in each diffusive state, *n* is the number of dimensions used to calculate *x*, and *Δt* is the length of time between two consecutive frames. To build the HMM, we installed and used the software vbSPT, version 1.1 on MATLAB version 2014a (17). Cells were rotated such that the long cell-axis corresponded to the x-axis. We used only consecutive frame trajectories and only displacements in the x-direction for analysis.

### Determination of diffusive domain size

To determine the diffusive domain size, we first calculated the mean squared displacement from all trajectories. To prevent bias from longer trajectories, we calculated the mean squared displacement for each timelag possible and used those mean values to calculate the total mean squared displacement from all trajectories. We then fit the MSD to an equation determined from the diffusion equation assuming a boundary along a single dimension of size L:

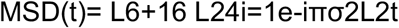

where *L* is the length of the boundary and σ is a value related to the diffusion coefficient by the relation *D* = *σ*^2^/2*n*, where n is the dimensions used to calculate the MSD. We solved this solution numerically by performing least squares minimization. Error in the fits were determined through a pseudo-bootstrapping method by resampling the trajectories used to calculate the MSD and determining the best fit parameters 100 times, and taking the standard deviation of these values.

### Growth rate comparison between rpoC-PAmCherry and MG1655

To assess the growth and functionality of the rpoC-PAmCherry fusion strain, we compared the growth of the MG1655 parental strain to our rpoC-PAmCherry fusion strain at 24°C. Five cultures from distinct isolated colonies for each strain were inoculated in EZRDM media without antibiotics. Cells were shaken overnight at room temperature (~ 16 hours). Saturated cultures were diluted back 1:200 and the OD at 600nm was monitored over time. Three technical replicates of measurements for each timepoint were taken for each biological replicate. Data was pooled and averaged over all biological and technical replicates and the doubling time was determined by fitting to an exponential equation.

### Western blot of RNAP

To assess whether the rpoC-PAmCherry protein was full-length and not proteolytically cleaved, we prepared cell lysates from an equivalent OD of cells for strains MG1655, MG1655:: rpoC-PAmCherry, and MG1655//pCH-FtsZ-mCherry grown as described above. Cell lysates were run on a 10% TBE protein gel (BioRad) and transferred at constant 25V for 2 hours at 4°C to a nitrocellulose membrane using a semi-dry transfer apparatus (GE). To ensure equal transfer, membranes were first blotted with Ponceau S stain to assess the transfer quality and rinsed with deionized water to remove the stain. Membrane blots were then incubated with 1x TBS + 0.1% Tween-20 solution (TBST) containing 5% w/v dry milk to block non-specific binding of the antibody for one hour at RT with gentle agitation, then washed three times for 5 minutes in TBST. Blocked membranes were then incubated with TBST + 1% w/v BSA (Sigma-Aldrich) with either a 1:50,000 dilution of anti-RpoC antibody (Biolegend clone NT73) or a 1:10,000 dilution of anti-mCherry antibody (Abcam ab167453) for one hour at RT with gentle agitation, then washed three times for 5 minutes in TBST. Membranes were then incubated in TBST with either 1:33,000 dilution of GAM-HRP secondary antibody (ThermoFisher 62-6520) for the anti-RpoC condition or GAR-HRP secondary antibody (BioRad #1705046) for the anti-mChery condition for one hour with gentle agitation, then washed five times for 5 minutes each in TBST. Membranes were then incubated with ECL solution diluted 1:5 in deionized water (BioRad #1705061) for 30 seconds, then wrapped in saran wrap and imaged using autoradiography film (Stellar Scientific BLI-810-100).

### Co-immunoprecipitation of rpoC(I1309A)-PAmCherry with rpoB

Cell lysates were prepared as described in Secition IV.G for strains MG1655, MG1655//pCH-PAmCherry, and MG1655//pCH-rpoC(I1309A)-PAmCherry. Cell lysates were incubated with Protein G sepharose beads (Sigma-Aldrich P3296) with anti-RpoC antibody (Biolegend clone NT73) overnight at 4°C. Bead incubated lysates were then washed with wash buffer (20 mM Tris–HCl pH 7.9, 150 mM KCl, 1 mM MgCl_2_) four times then incubated with 2x SDS sample buffer (1M Tris-HCl, pH 6.8, 50% glycerol, 10% SDS, 0.5% bromophenol blue, 0.5% β-mercaptoethanol) and boiled at 95% to elute protein. Samples at each stage were collected and run on a 10% TBE gel and transferred and detected as decribed in Section IV.G, but with detection using anti-RpoB antibody (BioLegend NT63) instead of anti-RpoC.

### DNA staining and image collection

For DNA nucleoid imaging experiments, cells were incubated with Hoechst 33342 dye (bisbenzimide H33342 trihydrochloride, ThermoFisher H3570) at a concentration of 10 μg/mL for fifteen minutes at room temperature with cell agitation; cell culture tubes were wrapped in aluminum foil to prevent the dye from bleaching. Cells were then fixed in 3% paraformaldehyde in a PBS solution for 15 minutes at room temperature. Cells were then washed twice with PBS. Fixed and stained cells were mixed 1:1 with anti-fading solution (20% n-propyl gallate, 60% glycerol, 1x PBS) and adhered to #1.5 coverslips treated with 0.01% poly-lysine solution. Coverslips were sealed onto an imaging coverglass using clear nail polish and imaged on a GE OMX SR structured illumination microscope (excitation: 405, camera channel: 488, exposure 50ms, 5% of highest laser intensity). Images were reconstructed using the standard parameters for the GE OMX SR microscope within the GE SRx software.

### Model simulations

To test the accuracy of output HMM parameters as a function of trajectory length and total number of displacements, simulations were performed. Trajectories were simulated within a 4 x 1.5 micron cell, modeled as a 2.5 x 1.5 micron cylinder with two half spherical ends with radius of 0.75 microns. Trajectory lengths were taken from a random exponential distribution with average trajectory length from 1 to 10 frames. The simulations were done with two diffusive states of 0.1 μm^2^/s and 1.0 μm^2^/s and simulated with timestep of 5ms, comparable to our imaging conditions. We simulated enough trajectories for each average trajectory length to have over 500,000 total displacements. From this set, we randomly selected 20, 50, 75, and 150 x 103 displacements to use for HMM analysis. These simulations were performed for two sets of kinetic transitions, k = 5 s^-1^ and k = 15 s^-1^; the rates from both diffusive states were the same in each condition (such that both would have occupancy of 50% each). We then compared the output parameters solved from the HMM to our input ground truth, which is plotted in Figure S13A and B.

To test whether having a “hidden” state that has the same diffusion coefficient as another state but significantly different transition kinetics, we again performed simulations of single particle trajectories within a 4 x 1.5 micron cell with a timestep of 5 ms. We simulated trajectories based off a model with three diffusive states, two with a diffusion coefficient of 0.1 μm2/s and one with 1.0 μm2/s and state occupancies of 10%, 45% and 45% respectively, hereto referred to as State 1a, State 1b, and State 2 respectively. We allowed transitions to occur between State 1b and State 2, with equal transition rates of 0.082 (or k ~ 13.3 s-1) while disallowing transitions to State 1a. When we input these resulting trajectories to solve the HMM, we found that the HMM output two states, with diffusion coefficients of 0.1 μm2/s and 1.0 μm2/s and occupancies approximately equal to the sum of the States 1a and 1b for the 0.1 μm2/s diffusive state. As would be predicted by steady state dynamics and the capacity of HMMs, we found that the transition rates were the weighted average of the two ground truth transition rates. This would imply that our observation of non-steady state dynamics is not a result of a “hidden” state.

## Notes

### Competing Interest Statement

The authors have declared no competing interest.

## References

Afek, A., Schipper, J. L., Horton, J., Gordân, R., & Lukatsky, D. B. (2014). Protein-DNA binding in the absence of specific base-pair recognition. Proceedings of the National Academy of Sciences, 111(48), 17140–17145. https://doi.org/10.1073/pnas.1410569111

Bakshi, S., Choi, H., Mondal, J., & Weisshaar, J. C. (2014). Time-dependent effects of transcription- and translation-halting drugs on the spatial distributions of the Escherichia coli chromosome and ribosomes. Molecular Microbiology, 94(4), 871–887. https://doi.org/10.1111/mmi.12805

Bakshi, S., Dalrymple, R. M., Li, W., Choi, H., & Weisshaar, J. C. (2013). Partitioning of RNA Polymerase Activity in Live Escherichia coli from Analysis of Single-Molecule Diffusive Trajectories. Biophysical Journal, 105(12), 2676–2686. https://doi.org/10.1016/j.bpj.2013.10.024

Bakshi, S., Siryaporn, A., Goulian, M., & Weisshaar, J. C. (2012). Superresolution imaging of ribosomes and RNA polymerase in live Escherichia coli cells. Molecular Microbiology, 85(1), 21–38. https://doi.org/10.1111/j.1365-2958.2012.08081.x

Bauer, M., & Metzler, R. (2012). Generalized Facilitated Diffusion Model for DNA-Binding Proteins with Search and Recognition States. Biophysical Journal, 102(10), 2321–2330. https://doi.org/10.1016/j.bpj.2012.04.008

Bettridge, K., Verma, S., Weng, X., Adhya, S., & Xiao, J. (2021). Single-molecule tracking reveals that the nucleoid-associated protein HU plays a dual role in maintaining proper nucleoid volume through differential interactions with chromosomal DNA. Molecular Microbiology, 115(1), 12–27. https://doi.org/10.1111/mmi.14572

Bhatia, R. P., Kirit, H. A., Predeus, A. v., & Bollback, J. P. (2022). Transcriptomic profiling of Escherichia coli K-12 in response to a compendium of stressors. Scientific Reports, 12(1), 8788. https://doi.org/10.1038/s41598-022-12463-3

Bohrer, C. H., & Xiao, J. (2020). Complex Diffusion in Bacteria. Advances in Experimental Medicine and Biology, 1267, 15–43. https://doi.org/10.1007/978-3-030-46886-6_2

Borukhov, S., & Nudler, E. (2008). RNA polymerase: the vehicle of transcription. Trends in Microbiology, 16(3), 126–134. https://doi.org/10.1016/j.tim.2007.12.006

Cabrera, J. E., Cagliero, C., Quan, S., Squires, C. L., & Jin, D. J. (2009). Active Transcription of rRNA Operons Condenses the Nucleoid in Escherichia coli: Examining the Effect of Transcription on Nucleoid Structure in the Absence of Transertion. Journal of Bacteriology, 191(13), 4180–4185. https://doi.org/10.1128/JB.01707-08

Cabrera, J. E., & Jin, D. J. (2003). The distribution of RNA polymerase in Escherichia coli is dynamic and sensitive to environmental cues. Molecular Microbiology, 50(5), 1493–1505. https://doi.org/10.1046/j.1365-2958.2003.03805.x

Campbell, E. A., Korzheva, N., Mustaev, A., Murakami, K., Nair, S., Goldfarb, A., & Darst, S. A. (2001). Structural mechanism for rifampicin inhibition of bacterial rna polymerase. Cell, 104(6), 901–912. https://doi.org/10.1016/s0092-8674(01)00286-0

Chen, J., Wassarman, K. M., Feng, S., Leon, K., Feklistov, A., Winkelman, J. T., Li, Z., Walz, T., Campbell, E. A., & Darst, S. A. (2017). 6S RNA Mimics B-Form DNA to Regulate Escherichia coli RNA Polymerase. Molecular Cell, 68(2), 388–397.e6. https://doi.org/10.1016/j.molcel.2017.09.006

Coltharp C., Buss, J., Plumer, T., Xiao, J. (2016). Proceedings of the National Academy of Sciences, 113(8), E1044–53. doi: 10.1073/pnas.1514296113.

Covert, M. W., & Palsson, B. Ø. (2002). Transcriptional Regulation in Constraints-based Metabolic Models of Escherichia coli. Journal of Biological Chemistry, 277(31), 28058–28064. https://doi.org/10.1074/jbc.M201691200

Dame, R. T., Rashid, F.-Z. M., & Grainger, D. C. (2020). Chromosome organization in bacteria: mechanistic insights into genome structure and function. Nature Reviews Genetics, 21(4), 227–242. https://doi.org/10.1038/s41576-019-0185-4

Dennis, P. P. (1976). Effects of chloramphenicol on the transcriptional activities of ribosomal RNA and ribosomal protein genes in Escherichia coli. Journal of Molecular Biology, 108(3), 535–546. https://doi.org/10.1016/S0022-2836(76)80135-0

Elowitz, M. B., Surette, M. G., Wolf, P.-E., Stock, J. B., & Leibler, S. (1999). Protein Mobility in the Cytoplasm of Escherichia coli. Journal of Bacteriology, 181(1), 197–203. https://doi.org/10.1128/JB.181.1.197-203.1999

Fazal, F. M., Meng, C. A., Murakami, K., Kornberg, R. D., & Block, S. M. (2015). Real-time observation of the initiation of RNA polymerase II transcription. Nature, 525(7568), 274–277. https://doi.org/10.1038/nature14882

Feklistov, A. (2013). RNA polymerase: in search of promoters. Annals of the New York Academy of Sciences, 1293(1), 25–32. https://doi.org/10.1111/nyas.12197

Feric, M., & Misteli, T. (2021). Phase separation in genome organization across evolution. Trends in Cell Biology, 31(8), 671–685. https://doi.org/10.1016/j.tcb.2021.03.001

Geffroy, L., Brown, H. A., DeVeaux, A. L., Koropatkin, N. M., & Biteen, J. S. (2022). Singlemolecule dynamics of surface lipoproteins in bacteroides indicate similarities and cooperativity. Biophysical Journal, 121(23), 4644–4655. https://doi.org/10.1016/j.bpj.2022.10.024

Gotta, S. L., Miller, O. L., & French, S. L. (1991). rRNA transcription rate in Escherichia coli. Journal of Bacteriology, 173(20), 6647–6649. https://doi.org/10.1128/jb.173.20.6647-6649.1991

Graham, M. Y., Tal, M., & Schlessinger, D. (1982). lac Transcription in Escherichia coli cells treated with chloramphenicol. Journal of Bacteriology, 151(1), 251–261. https://doi.org/10.1128/jb.151.1.251-261.1982

Hatoum, A., & Roberts, J. (2008). Prevalence of RNA polymerase stalling at Escherichia coli promoters after open complex formation. Molecular Microbiology, 6δ(1), 17–28. https://doi.org/10.1111/j.1365-2958.2008.06138.x

Henderson, K. L., Felth, L. C., Molzahn, C. M., Shkel, I., Wang, S., Chhabra, M., Ruff, E. F., Bieter, L., Kraft, J. E., & Record, M. T. (2017). Mechanism of transcription initiation and promoter escape by E. coli RNA polymerase. Proceedings of the National Academy of Sciences, 114(15). https://doi.org/10.1073/pnas.1618675114

Herring, C. D., Raffaelle, M., Allen, T. E., Kanin, E. I., Landick, R., Ansari, A. Z., & Palsson, B. Ø. (2005). Immobilization of Escherichia coli RNA polymerase and location of binding sites by use of chromatin immunoprecipitation and microarrays. Journal of Bacteriology, 187(17), 6166–6174. https://doi.org/10.1128/JB.187.17.6166-6174.2005

Hinton, D. M., Pande, S., Wais, N., Johnson, X. B., Vuthoori, M., Makela, A., & Hook-Barnard, I. (2005). Transcriptional takeover by sigma appropriation: remodelling of the sigma70 subunit of Escherichia coli RNA polymerase by the bacteriophage T4 activator MotA and co-activator AsiA. Microbiology (Reading, England), 151(Pt 6), 1729–1740. https://doi.org/10.1099/mic.0.27972-0

Jishage, M., Iwata, A., Ueda, S., & Ishihama, A. (1996). Regulation of RNA polymerase sigma subunit synthesis in Escherichia coli: intracellular levels of four species of sigma subunit under various growth conditions. Journal of Bacteriology, 178(18), 5447–5451. https://doi.org/10.1128/jb.178.18.5447-5451.1996

Ladouceur, A.-M., Parmar, B. S., Biedzinski, S., Wall, J., Tope, S. G., Cohn, D., Kim, A., Soubry, N., Reyes-Lamothe, R., & Weber, S. C. (2020). Clusters of bacterial RNA polymerase are biomolecular condensates that assemble through liquid-liquid phase separation. Proceedings of the National Academy of Sciences of the United States of America, 117(31), 18540–18549. https://doi.org/10.1073/pnas.2005019117

Li, G.-W., Burkhardt, D., Gross, C., & Weissman, J. S. (2014). Quantifying Absolute Protein Synthesis Rates Reveals Principles Underlying Allocation of Cellular Resources. Cell, 157(3), 624–635. https://doi.org/10.1016/j.cell.2014.02.033

Liao, Y., Li, Y., Schroeder, J. W., Simmons, L. A., & Biteen, J. S. (2016). Single-Molecule DNA Polymerase Dynamics at a Bacterial Replisome in Live Cells. Biophysical Journal, 111(12), 2562–2569. https://doi.org/10.1016/j.bpj.2016.11.006

Macvanin, M., & Adhya, S. (2012). Architectural organization in E. coli nucleoid. Biochimica et Biophysica Acta, 1819(7), 830–835. https://doi.org/10.1016/j.bbagrm.2012.02.012

Minton, A. P. (2001). The Influence of Macromolecular Crowding and Macromolecular Confinement on Biochemical Reactions in Physiological Media. Journal of Biological Chemistry, 276(14), 10577–10580. https://doi.org/10.1074/jbc.R100005200

Orsini, G., Ouhammouch, M., le Caer, J. P., & Brody, E. N. (1993). The asiA gene of bacteriophage T4 codes for the anti-sigma 70 protein. Journal of Bacteriology, 175(1), 85–93. https://doi.org/10.1128/jb.175.1.85-93.1993

Persson, F., Lindén, M., Unoson, C., & Elf, J. (2013). Extracting intracellular diffusive states and transition rates from single-molecule tracking data. Nature Methods, 10(3), 265–269. https://doi.org/10.1038/nmeth.2367

Sastry, A. v, Gao, Y., Szubin, R., Hefner, Y., Xu, S., Kim, D., Choudhary, K. S., Yang, L., King, Z. A., & Palsson, B. O. (2019). The Escherichia coli transcriptome mostly consists of independently regulated modules. Nature Communications, 10(1), 5536. https://doi.org/10.1038/s41467-019-13483-w

Seshasayee, A. S. N., Sivaraman, K., & Luscombe, N. M. (2011). An Overview of Prokaryotic Transcription Factors (pp. 7–23). https://doi.org/10.1007/978-90-481-9069-0_2

Severinova, E., Severinov, K., & Darst, S. A. (1998). Inhibition of Escherichia coli RNA polymerase by bacteriophage T4 AsiA. Journal of Molecular Biology, 279(1), 9–18. https://doi.org/10.1006/jmbi.1998.1742

Sharma, U. K., & Chatterji, D. (2008). Differential Mechanisms of Binding of Anti-Sigma Factors Escherichia coli Rsd and Bacteriophage T4 AsiA to E. coli RNA Polymerase Lead to Diverse Physiological Consequences. Journal of Bacteriology, 190(10), 3434–3443. https://doi.org/10.1128/JB.01792-07

Stracy, M., & Kapanidis, A. N. (2017). Single-molecule and super-resolution imaging of transcription in living bacteria. Methods, 120, 103–114. https://doi.org/10.1016/j.ymeth.2017.04.001

Stracy, M., Lesterlin, C., Garza de Leon, F., Uphoff, S., Zawadzki, P., & Kapanidis, A. N. (2015). Live-cell superresolution microscopy reveals the organization of RNA polymerase in the bacterial nucleoid. Proceedings of the National Academy of Sciences, 112(32). https://doi.org/10.1073/pnas.1507592112

Stracy, M., Schweizer, J., Sherratt, D. J., Kapanidis, A. N., Uphoff, S., & Lesterlin, C. (2021). Transient non-specific DNA binding dominates the target search of bacterial DNA-binding proteins. Molecular Cell, 81(7), 1499–1514.e6. https://doi.org/10.1016/j.molcel.2021.01.039

Taniguchi, Y., Choi, P. J., Li, G.-W., Chen, H., Babu, M., Hearn, J., Emili, A., & Xie, X. S. (2010). Quantifying E. coli Proteome and Transcriptome with Single-Molecule Sensitivity in Single Cells. Science, 329(5991), 533–538. https://doi.org/10.1126/science.1188308

Terry, B. R., Matthews, E. K., & Haseloff, J. (1995). Molecular Characterization of Recombinant Green Fluorescent Protein by Fluorescence Correlation Microscopy. Biochemical and Biophysical Research Communications, 217(1), 21–27. https://doi.org/10.1006/bbrc.1995.2740

Uphoff, S., Reyes-Lamothe, R., Garza de Leon, F., Sherratt, D. J., & Kapanidis, A. N. (2013). Single-molecule DNA repair in live bacteria. Proceedings of the National Academy of Sciences, 110(20), 8063–8068. https://doi.org/10.1073/pnas.1301804110

van Helvoort, J. M., Kool, J., & Woldringh, C. L. (1996). Chloramphenicol causes fusion of separated nucleoids in Escherichia coli K-12 cells and filaments. Journal of Bacteriology, 178(14), 4289–4293. https://doi.org/10.1128/jb.178.14.4289-4293.1996

Vo, N. v., Hsu, L. M., Kane, C. M., & Chamberlin, M. J. (2003). In Vitro Studies of Transcript Initiation by Escherichia coli RNA Polymerase. 3. Influences of Individual DNA Elements within the Promoter Recognition Region on Abortive Initiation and Promoter Escape. Biochemistry, 42(13), 3798–3811. https://doi.org/10.1021/bi026962v

Wang, F., Redding, S., Finkelstein, I. J., Gorman, J., Reichman, D. R., & Greene, E. C. (2013). The promoter-search mechanism of Escherichia coli RNA polymerase is dominated by three-dimensional diffusion. Nature Structural & Molecular Biology, 20(2), 174–181. https://doi.org/10.1038/nsmb.2472

Wang, G., Hauver, J., Thomas, Z., Darst, S. A., & Pertsinidis, A. (2016). Single-Molecule Real-Time 3D Imaging of the Transcription Cycle by Modulation Interferometry. Cell, 167(7), 1839–1852.e21. https://doi.org/10.1016/j.cell.2016.11.032

Weng, X., Bohrer, C. H., Bettridge, K., Lagda, A. C., Cagliero, C., Jin, D. J., & Xiao, J. (2019). Spatial organization of RNA polymerase and its relationship with transcription in Escherichia coli. Proceedings of the National Academy of Sciences, 116(40), 20115–20123. https://doi.org/10.1073/pnas.1903968116

Wu, C., Balakrishnan, R., Braniff, N., Mori, M., Manzanarez, G., Zhang, Z., & Hwa, T. (2022). Cellular perception of growth rate and the mechanistic origin of bacterial growth law. Proceedings of the National Academy of Sciences, 119(20). https://doi.org/10.1073/pnas.2201585119

Xiang, Y., Surovtsev, I. v., Chang, Y., Govers, S. K., Parry, B. R., Liu, J., & Jacobs-Wagner, C. (2021). Interconnecting solvent quality, transcription, and chromosome folding in Escherichia coli. Cell, 184(14), 3626–3642.e14. https://doi.org/10.1016/j.cell.2021.05.037

Yamazaki, K., Nagata, A., Kano, Y., & Imamoto, F. (1984). Isolation and characterization of nucleoid proteins from Escherichia coli. Molecular & General Genetics: MGG, 196(2), 217–224. https://doi.org/10.1007/BF00328053

Zaid, I. M., Lomholt, M. A., & Metzler, R. (2009). How Subdiffusion Changes the Kinetics of Binding to a Surface. Biophysical Journal, 97(3), 710–721. https://doi.org/10.1016/j.bpj.2009.05.022

Zhang, N., Schäfer, J., Sharma, A., Rayner, L., Zhang, X., Tuma, R., Stockley, P., & Buck, M. (2015). Mutations in RNA Polymerase Bridge Helix and Switch Regions Affect Active-Site Networks and Transcript-Assisted Hydrolysis. Journal of Molecular Biology, 427(22), 3516–3526. https://doi.org/10.1016/j.jmb.2015.09.005

Zimmerman, S. B. (2006). Shape and compaction of Escherichia coli nucleoids. Journal of Structural Biology, 156(2), 255–261. https://doi.org/10.1016/j.jsb.2006.03.022

