## SupplementalFigures for "RNAP promoter search and transcription kinetics in live *E. coli* cells"

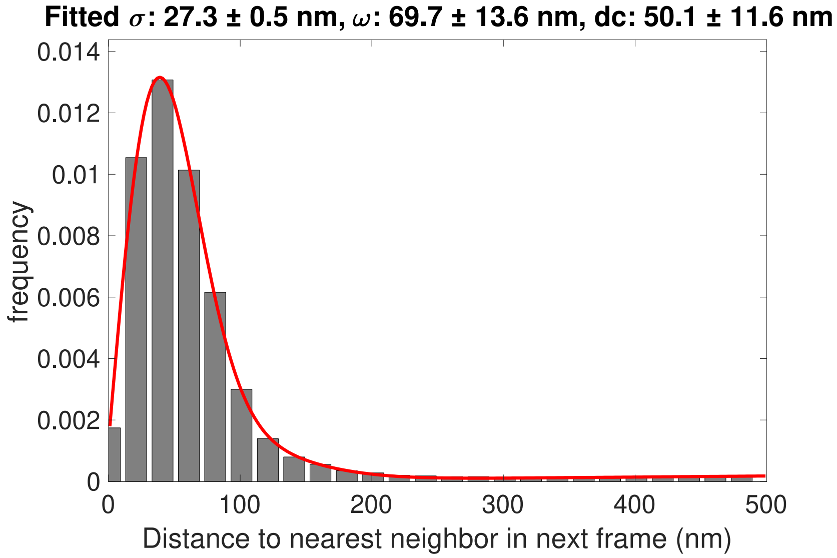


**Supplementary Figure 1: Calculation of the localization precision in single-molecule tracking (SMT) using the Nearest Neighbor Distance Distribution.** Cells expressing RpoC-PAmCherry were fixed and single RpoC-PAmCherry molecules were photoactivated and localized sparsely. Repeat localizations from same molecules were used to compute the nearest neighbor distances and pooled to construct the distribution histogram. The histogram was fit using the equation described in (Endesfelder et al., 2014) and (Coltharp et al., 2016) with fitting parameters shown above the graph.

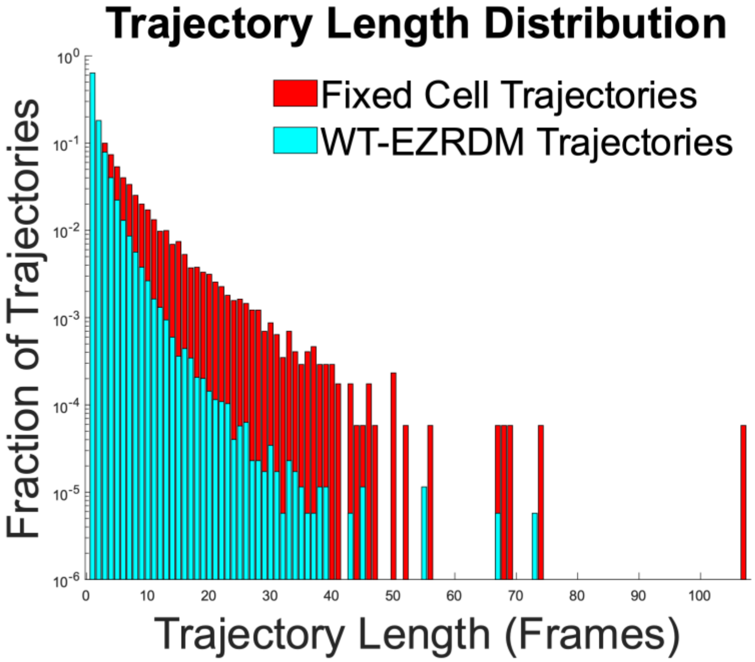


**Supplementary Figure 2: Live cell single-molecule tracking is not limited by photobleaching. Trajectory length in fixed cells and in WT-EZRDM cells.** The average trajectory length of PAmCherry molecules in fixed cells is $6.0 \pm 5.9$ frames (from 10,727 trajectories in 45 cells, red), longer than the average trajectory length for PAmCherry molecules in live cells (WT-EZRDM, $3.4 \pm 2.4$ frames, 63,182 trajectories from 353 cells, cyan). Single frame trajectories were excluded from the analysis.

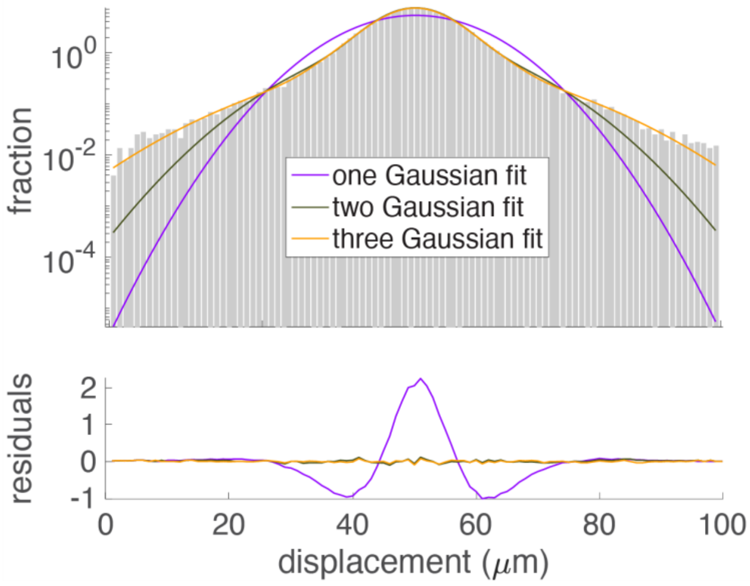


**Supplementary Figure 3: Single-step displacement distribution of RNAP-PAmCherry SMT trajectories** (gray bars, in log scale) in EZRDM growth medium shows that a three-population Gaussian fit (orange) best describes the distribution than single- (purple) and two-population (black) fits. Residuals for these fits are plotted below in linear scale.

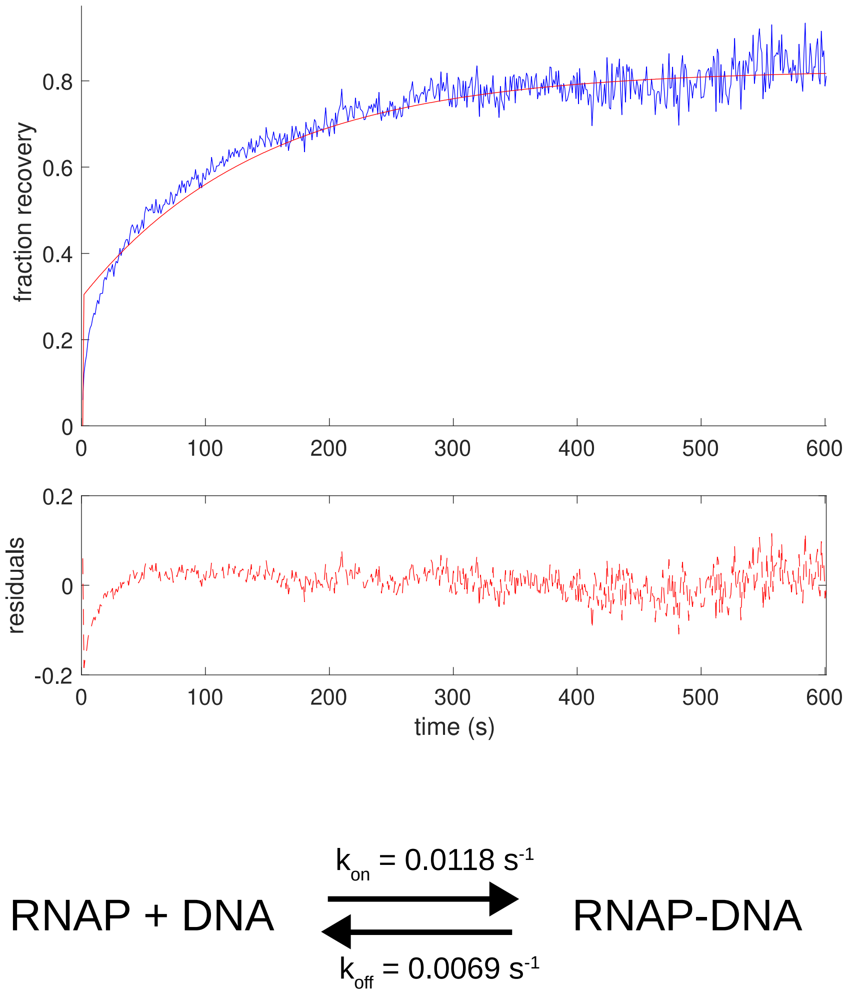


**Supplementary Figure 4: A one-step binding model without involving a RNAP-DNA elongation complex does not describe well the FRAP curve of RNAP-GFP.** The average FRAP recovery for RNAP-GFP is plotted in blue, with the red solid line indicating the FRAP curve resulting from the best fit one-step binding model. Residuals for this model against the data are plotted below and show significant deviations, particularly at the early timepoints.

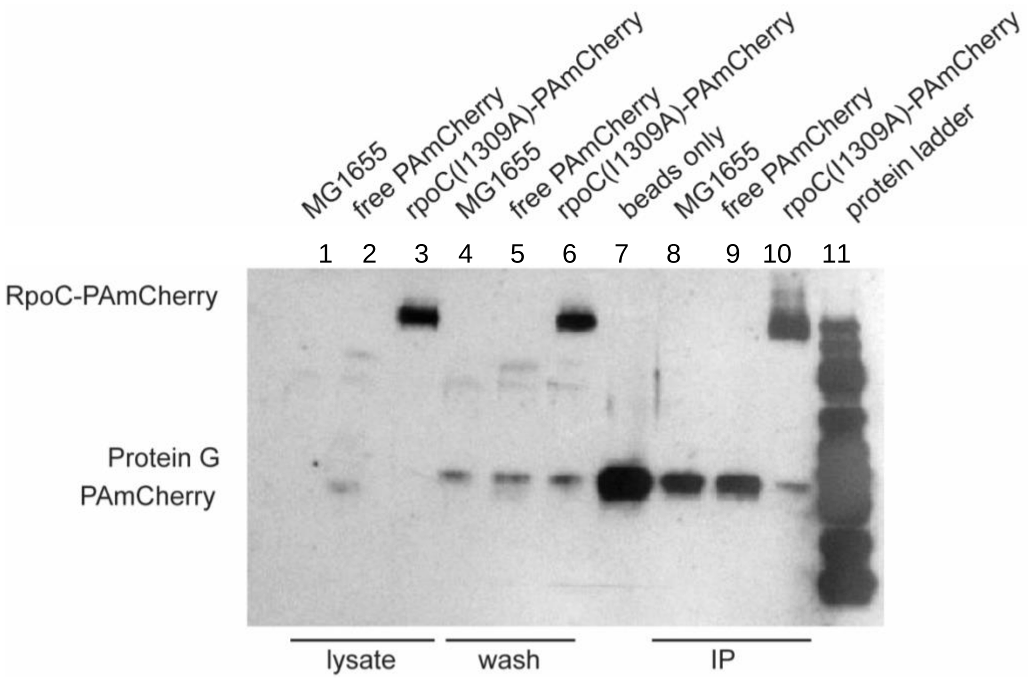


**Supplementary Figure 5: Coimmunoprecipitation of RpoC(I1309A)-PAmCherry demonstrates its incorporation into the holoenzyme.** Cell lysates from MG1655 (the parental strain), MG1655//pCH-PAmCherry, and MG1655//pCH-rpoC(I1309A)-PAmCherry were incubated with beads containing anti-RpoB antibody and detected on protein blots using anti-mCherry antibodies. Lanes 1-3 are the whole cell lysate, lanes 4-6 are the wash flow through, lane 7 is the beads-only control, and lanes 8-10 is the immunoprecipitation elute. The presence of the RpoC(I1309A)-PAmCherry protein in lane 10 indicates that this mutant RpoC subunit is incorporated into the holoenzyme.

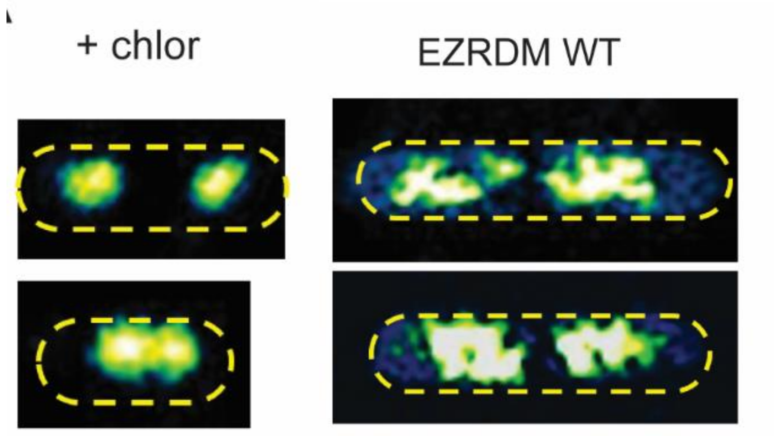


**Supplementary Figure 6: Nucleoid compaction induced by chloramphenicol.** Representative images showing chloramphenicol-induced nucleoid compaction compared to WT. The chromosomal DNA was stained with Hoechst 33342 dye (10 μg/mL, 15 minutes).

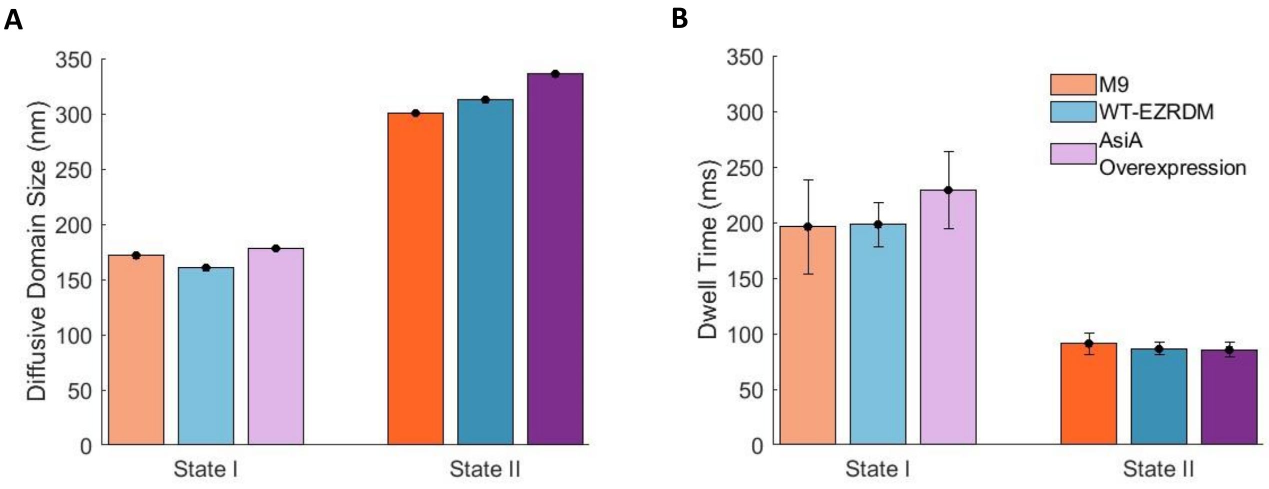


**Supplementary Figure 7: Diffusive domain size and dwell time are similar among M9, AsiA Overexpression, and WT.** Average diffusive domain size for state I and state II RNAP molecules for M9 (orange) and AsiA Overexpression (purple) conditions in comparison with WT condition (cyan). Domain size were calculated using the Kusumi equation. **B**. Average dwell time comparison for state I and state II RNAP molecules for M9 (orange) and AsiA Overexpression (purple) conditions in comparison with WT condition (cyan). Error bars represent standard error on the mean for both **A** and **B**.

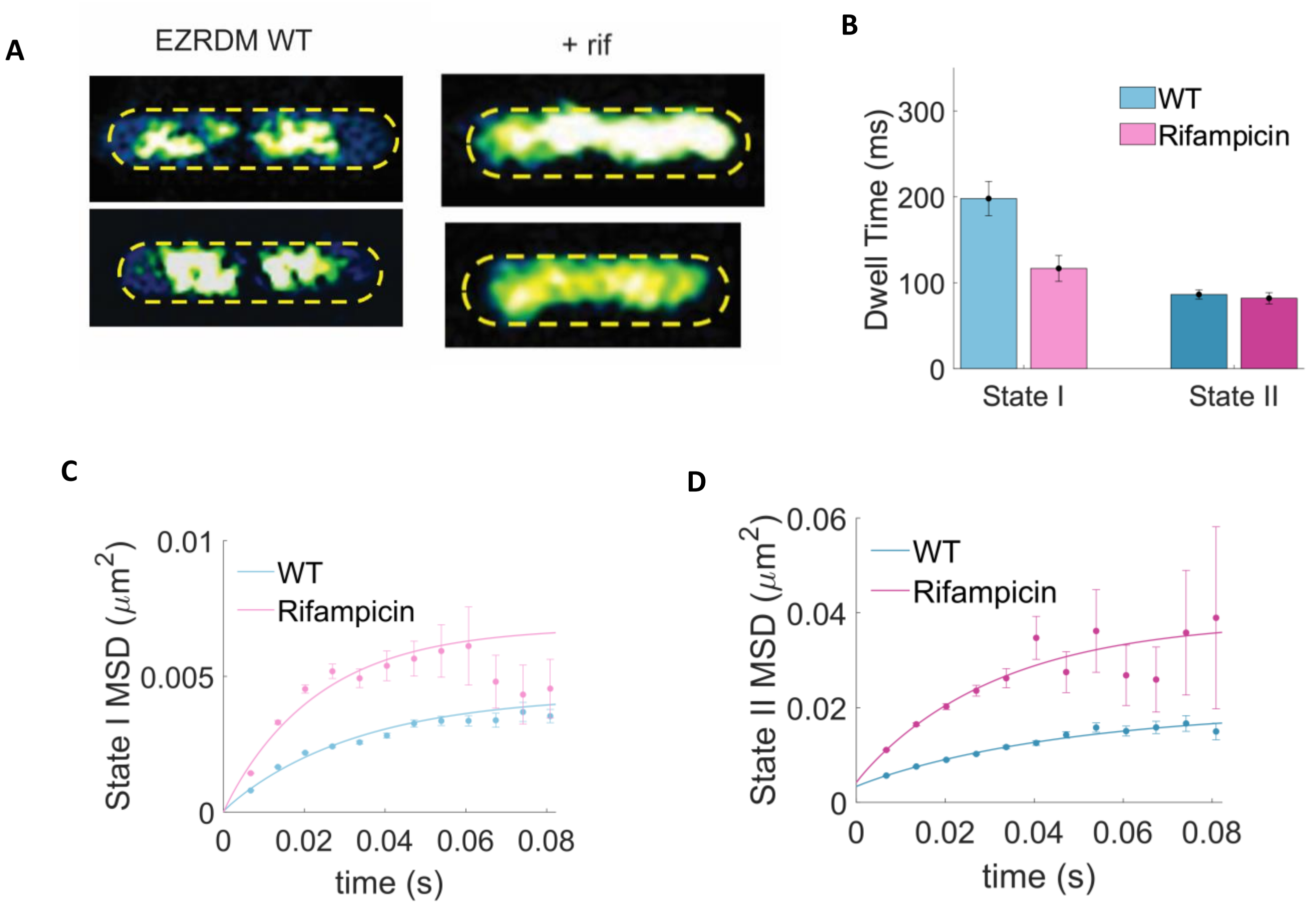


**Supplementary Figure 8: Rifampicin causes expanded nucleoid and diffusive behavior. A**. Representative images showing rifampicin-induced nucleoid expansion compared to WT. DNA is stained with Hoechst 33342 dye (10 μg/mL,15 minutes). **B**. Average dwell time for state I and state II RNAP molecules for rifampicin-treated cells (pink) in comparison with WT cells (cyan). **C** and **D**. MSD plots for state I (**C**) and state II (**D**) RNAP molecules (pink) in comparison with WT condition (cyan).

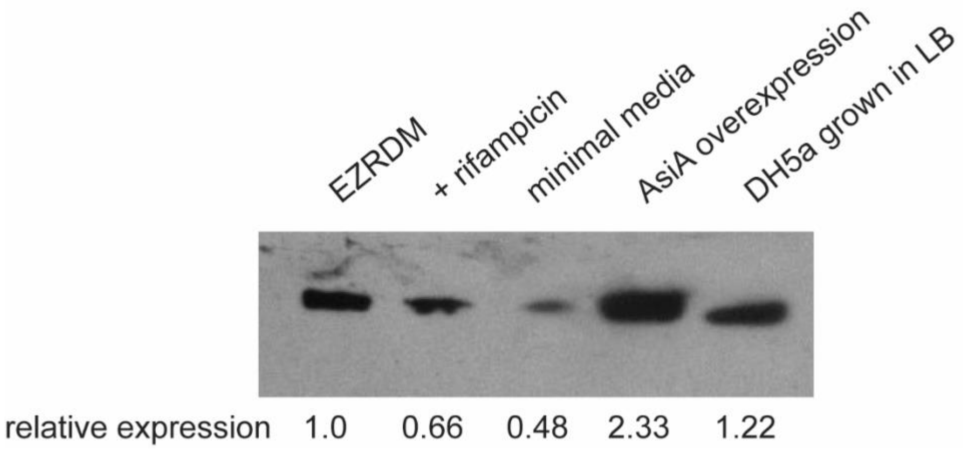


**Supplementary Figure 9: Relative expression levels of RNAP-PAmCherry under conditions used in this study.** Equal numbers of cells from each condition were boiled and run on a 10% gel, then probed with anti-RpoC antibody. Results were quantified using ImageJ and normalized to the EZRDM condition.

**Stokes-Einstein Calculation:**

$$D = \frac{k_B T}{6\pi\eta r}$$

$$D \propto \frac{1}{r}$$

$$D_{GFP} r_{GFP} = D_{RNAP} r_{RNAP}$$

$$7.7\ \mu m^2/s\ (2.8\ nm) = D_{RNAP}\ (10nm)$$

$$D_{RNAP} = 2.16\ \mu m^2/s$$

**On Rate and $K_d$ Calculation:**

We assigned state I RNAP-PAmCherry molecules to be bound to the DNA, and state II RNAP-PAmCherry molecules to be diffusing in the nucleoid. The binding reaction then following the equation below, with the apparent $k_{on}$ and $k_{off}$ of single RNAP molecules determined by HMM of SMT data.

$$RNAP + DNA \leftrightarrow RNAP \cdot DNA$$

Assuming that (1) the *E. coli* RNAP binds to a minimal 20-bp sequence, (2) each base pair of the chromosomal DNA contributes to a unique nonspecific binding site for RNAP, and (3) on average, there is one full chromosomes per *E. coli* cell under our growth medium condition, (4) an average *E. coli* cell has a size of 1 μm x 1 μm x 3 μm, we arrive at a concentration of chromosomal DNA binding sites at ~ 3.3 mM.

Assuming that RNAP is expressed at approximately 4000 molecules per cell, the cellular concentration of RNAP is 3 μM.

Since [DNA] >> [RNAP], we can use a first order approximation, where

$$K_d = \frac{k_{off}[DNA]}{k_{on}}$$

Therefore, using $k_{on} = k_{21} = 7.48\ s^{-1}$, and $k_{off} = k_{12} = 4.37\ s^{-1}$, the apparent $K_d$ for RNAP's nonspecific binding is calculated at ~ 2 mM.

**Table 1. vbSPT HMM Analysis of SMT Data**

| | **State I** | | | | **State II** | | | | **State III** | | | | | |
|---|---|---|---|---|---|---|---|---|---|---|---|---|---|---|
| | D1 (∝m$^2$/s) | Occupancy (%) | Dwell Time (ms) | Transition Rates ($s^{-1}$) | D2 (∝m$^2$/s) | Occupancy (%) | Dwell Time (ms) | Transition Rates ($s^{-1}$) | D3 (∝m$^2$/s) | Occupancy (%) | Dwell Time (ms) | Transition Rates ($s^{-1}$) | n (Traj.) | n (Cells) |
| WT-EZRDM | 0.10 ±0.002 | 43.5 ±1.2 | 198.0 ±20.0 | $k_{12}$ = 4.37 ±0.59 $k_{13}$ = 0.89 ±0.24 | 0.30 ±0.007 | 43.7 ±1.1 | 86.3 ±5.5 | $k_{21}$ = 7.48 ±0.76 $k_{23}$ = 5.29 ±0.55 | 1.4 ±0.03 | 12.8 ±0.4 | 28.6 ±1.4 | $k_{31}$ = 5.03 ±1.17 $k_{32}$ = 35.48 ±2.41 | 63,183 | 353 |
| RNAP I1309A | 0.10 ±0.003 | 49.9 ±1.5 | 267.8 ±75.2 | $k_{12}$ = 4.04 ±0.88 | 0.38 ±0.007 | 50.1 ±1.5 | 55.5 ±5.7 | $k_{21}$ = 19.48 ±1.93 | | | | | 13,473 | 364 |
| Chloram-phenicol | 0.092 ±0.001 | 48.1 ±0.7 | 278.3 ±28.9 | $k_{12}$ = 3.86 ±0.38 | 0.39 ±0.003 | 51.9 ±0.7 | 61.5 ±3.0 | $k_{21}$ = 17.45 ±0.82 | | | | | 44,168 | 121 |
| M9 | 0.11 ±0.003 | 44.2 ±1.3 | 196.0 ±42.3 | $k_{12}$ = 5.40 ±0.96 | 0.38 ±0.005 | 55.8 ±1.3 | 90.9 ±9.8 | $k_{21}$ = 11.64 ±1.18 | | | | | 21,500 | 127 |
| AsiA Overexpr. | 0.11 ±0.002 | 44.0 ±1.0 | 229.2 ±34.7 | $k_{12}$ = 4.62 ±0.60 | 0.39 ±0.004 | 56.0 ±1.0 | 85.6 ±6.7 | $k_{21}$ = 12.37 ±0.93 | | | | | 33,965 | 104 |
| Rifampicin | 0.17 ±0.004 | 36.7 ±0.8 | 116.7 ±15.1 | $k_{12}$ = 9.23 ±1.09 | 0.83 ±0.008 | 63.3 ±0.8 | 82.0 ±6.5 | $k_{21}$ = 13.14 ±1.00 | | | | | 40,144 | 361 |

$k_{12}$ means from State I to State II
Error bars represent standard error on the mean given by vbSPT

**Table 2. Confinement Analysis of SMT Data**

| | State I | State II | State III |
|---|---|---|---|
| | Confinement Zone Size, nm | Confinement Zone Size, nm | Confinement Zone Size, nm |
| WT-EZRDM | 160.5 ±0.08 | 312.6 ±0.3 | 337.1 ± 1.1 |
| RNAP I1309A | 248.3 ±0.2 | 533.8 ±0.5 | |
| Chloramphenicol | 141.8 ±0.004 | 297.1 ±0.2 | |
| M9 | 171.6 ±0.05 | 300.5 ±0.01 | |
| AsiA Overexpression | 178.1 ±0.02 | 336.0 ±0.1 | |
| Rifampicin | 202.1 ±0.2 | 453.2 ±0.2 | |

Errors are standard error on the mean.

**Table 3: Kinetic Modeling of FRAP Data**

| | $k_{on}$ ($s^{-1}$) | $k_{off}$ ($s^{-1}$) | $k_{esc}$ ($s^{-1}$) | $k_{term}$ ($s^{-1}$) | $P_{free}$ | $P_{bound}$ | $P_{transcribing}$ | Number of Cells |
|---|---|---|---|---|---|---|---|---|
| WT-EZRDM | 0.89 | 1.1 | 0.012 | 0.0083 | 33.9% | 26.5% | 39.6% | 45 |
| M9 | 0.63 | 1.36 | 0.019 | 0.0092 | 41.9% | 18.7% | 39.3% | 46 |
| AsiA Overexpression | 0.83 | 1.17 | 0.0088 | 0.0097 | 43.0% | 29.6% | 27.4% | 45 |
| Rifampicin | 0.0017 | 0.0011 | | | 61.0% | 38.9% | | 45 |
